# Ionic regulation of Gram-positive phage adsorption governs host range and improves phage isolation efficiency

**DOI:** 10.64898/2026.03.09.706807

**Authors:** Braira Wahid, Jinxin Zhao, Adam Truskewycz, Melanie MacGregor, Jack Ramsay, Morgyn Warner, Peter Speck

**Affiliations:** Bacteriophage Research and Innovation Laboratory (BRIL), College for Science and Engineering, Flinders University, Bedford Park 5042, South Australia; Department of Microbiology, Biomedicine Discovery Institute, Monash University, Melbourne, Victoria 3800, Australia; Institute for Nanoscale Science and Technology, College for Science and Engineering, Flinders University, Bedford Park 5042, South Australia; Faculty of Health and Medical Sciences, University of Adelaide, Adelaide, SA

**Keywords:** phage therapy, antimicrobial resistance, multidrug-resistance, Staphylococcus species, Gram-positive phages

## Abstract

Antimicrobial-resistant *Staphylococcus aureus* and *S. epidermidis* remain major causes of invasive infection, yet isolation of therapeutically robust lytic phages targeting Gram-positive pathogens is often constrained by adsorption barriers imposed by thick peptidoglycan cell walls. We demonstrate that these limitations are primarily methodological. By optimising the ionic microenvironment during isolation through divalent cation supplementation (10 mM CaCl_2_ and MgSO_4_), controlled enrichment, phage polyclonal mixtures, and limited lysozyme-assisted release, we recovered 28 genomically distinct lytic phages compared with six obtained using conventional protocols. Host-range profiling across 103 clinical isolates showed broad infectivity, with 51.7% of interactions supporting high-efficiency infection and multiple phages exhibiting cross-species activity. Ion-supplemented adaptive passaging restored and expanded lytic capacity against resistant strains within five rounds. These findings show that ionic regulation of adsorption reshapes Gram-positive phage-host dynamics and provide a scalable framework for precision targeting of resistant *Staphylococcus* infections.

## 1. Introduction

Bacteriophages (phages) are the most abundant biological entities on earth. They are capable of infecting bacteria and play a vital role in shaping microbial ecology and evolution [1]. Phage infection is a highly specific process that begins with adsorption to host cell surface receptors. This is typically followed by genome injection and replication, ultimately leading to cell lysis in virulent infections [2]. The adsorption stage is mediated by phage-encoded receptor binding proteins (RBPs) that interact with defined cell surface structures, making this step a primary determinant of host specificity and infection efficiency [3]. In phages that infect Gram-positive bacteria, these interactions often involve cell wall glycopolymers such as wall teichoic acids (WTAs) and lipoteichoic acids (LTAs), which serve as primary or secondary receptor sites for irreversible binding by the phage tail apparatus [4, 5].

Globally, Gram-positive bacteria are a significant and expanding cause of antimicrobial resistance (AMR), posing critical challenges for healthcare policy and infection control [6]. Strong antimicrobial stewardship and infection prevention measures are required due to the increasing prevalence of pathogens such as methicillin-susceptible *Staphylococcus aureus* (MSSA), methicillin-resistant *S. aureus* (MRSA), and multidrug-resistant (MDR) *S. aureus, Enterococcus faecium*, and Streptococcus spp. [7-9]. In particular, Gram-positive bacteria are major causative agents of bloodstream infections, wound infections, surgical device-associated infections, and other healthcare-associated infections. The fatality rates for invasive diseases caused by these bacteria range from 10% to 30%, highlighting the importance of targeted policy interventions [10].

Although there has been extensive research on phages, much of our understanding of their mechanisms still relies on their infection dynamics in Gram-negative hosts. The outer membrane’s lipopolysaccharides (LPS) and protein receptors of these bacteria have been studied in detail [11]. In contrast to this, studies on Gram-positive bacteriophages have been relatively underrepresented because of the structural complexity of the thick, peptidoglycan-rich cell envelope and its associated polymers, which produce a distinct repertoire of receptors and unique barriers to phage adsorption and propagation [12]. Recent advancements in genomics and structural biology have begun to illustrate the role of teichoic acid and other cell wall components in the interactions between Gram-positive bacteria and phages [13]. Isolating phages targeting Gram-positive bacteria remains a significant challenge and requires urgent attention. Unlike Gram-negative bacteria, whose lipopolysaccharide (LPS) and outer membrane facilitate phage binding and enrichment using standard isolation protocols, Gram-positive bacteria possess a thick peptidoglycan cell wall that restricts phage adsorption and reduces recovery efficiency under conventional enrichment conditions [14, 15].

The limited development of phage therapeutics targeting key Gram-positive pathogens such as MDR *S. aureus* reflects persistent methodological barriers to phage isolation and propagation. The problem is exacerbated by the emergence of vancomycin-resistant Gram-positive strains, for which treatment options are increasingly limited, underscoring the urgency of developing efficient phage isolation and optimisation strategies. In Australia, this challenge has direct clinical implications: patients requiring compassionate phage therapy in South Australia currently depend on interstate phage banks, where strain-specific mismatches and logistical delays can preclude timely treatment. The growing burden of invasive MRSA infections, including bacteraemia, highlights the urgent need for locally responsive phage discovery platforms aligned with regional epidemiology.

In contrast to Gram-negative systems, the recovery of lytic phages from Gram-positive bacteria often requires specialised approaches that overcome the structural limitations imposed by the peptidoglycan layer. While adding divalent cations to growth media improves electrostatic interactions necessary for effective phage adsorption, chemical or enzymatic alteration of the cell wall can temporarily expose otherwise inaccessible receptors [16]. Likewise, phage susceptibility in *S. epidermidis* is governed by complex and highly variable cell wall teichoic acid structures and diverse anti-phage defence systems, and comparatively few lytic *S. epidermidis* phages have been characterised to date [17]. We hypothesised that the limited recovery of phages from Gram-positive bacteria under standard isolation protocols is driven by suboptimal adsorption conditions linked to host physiological state and ionic environment. Based on this, we developed a modified isolation framework optimised for Gram-positive hosts that integrates (i) mid-log phase host cultivation to maximise receptor availability, (ii) targeted manipulation of divalent ion concentrations to enhance adsorption kinetics, and (iii) strategic use of cell wall-modifying agents to facilitate phage access. Compared with published SOPs [18-20], our optimised isolation framework substantially increased recovery efficiency, yielding 28 distinct phages (24 *S. aureus* and 4 *S. epidermidis*) from a batch of 56 host strains.

## 2. Methods

### 2.1. Bacterial strains

A panel of more than 50 *Staphylococcus aureus* isolates, including MSSA, MRSA, and multidrug-resistant (MDR) strains representing 15 distinct sequence types, was collected from a local diagnostic laboratory, with most strains originating from infected wounds or diabetic foot ulcers. The antibiograms and sequence types of these strains are listed in **Supplementary File S1**. The host range of the isolated phages was tested against a total of 103 *S. aureus* and *S. epidermidis* strains.

### 2.2. Phage isolation

A mixture of raw sewage samples collected from three different locations (add locations) was initially processed for phage isolation. We initially performed phage isolation using the conventional Phage-on-Tap protocol [18], which is highly efficient for Gram-negative hosts. However, only 6 phages were recovered from 56 Gram-positive strains. We therefore optimised the enrichment strategy. In the modified approach, 3-5 mL of sewage samples was combined with 5 mL of 2X tryptic soy broth (TSB) supplemented with 10 mM MgSO_4_ and CaCl_2_ to enhance adsorption and recovery efficiency. After adding 100-200 µL of mid-log-phase bacterial culture for enrichment, the mixtures were incubated in a shaking incubator at 37 °C for 4-6 hours, depending on the appearance of turbidity. Unlike conventional phage isolation methods that often rely on overnight enrichment, we employed controlled, shorter incubation periods to minimise selective pressures that can favour phage-resistant subpopulations or rapidly replicating narrow host-range variants. After incubation, the cultures were filtered through syringe filters (0.45 µm pore size, 25 mm PVDF (Polyvinylidene fluoride), Sigma Aldrich) to remove larger debris and bacterial cells. Chloroform (1.2% (v/v)) was added to the resulting filtrates to inactivate any remaining bacteria and release the intracellular phage particles. After chloroform treatment, samples were centrifuged to remove insoluble material and residual organic phase, and the clarified supernatants were collected as crude phage lysates using 0.22 µm pore size syringe filter. For some challenging and phage-resistant strains, a low dose of lysozyme (≤10 µg/mL) was used to achieve partial cell wall rupture and improved phage recovery; this step was employed selectively on some isolates only. To maximise recovery and long-term usability of *Staphylococcus*-infecting phages, all raw phage lysates obtained during routine screening were stored regardless of their immediate lytic activity against the corresponding host strain. These lysates were pooled to generate a composite, polyclonal phage reservoir, which was used for parallel host-specific enrichment in our routine phage discovery work against newly obtained *Staphylococcus* isolates. This strategy eliminates the need for repeated environmental sampling (e.g., sewage collection) while enabling rapid screening of a polyclonal phage mixture against clinical isolates, followed by phage purification and amplification. This pooling preserves low-abundance and genetically diverse phage variants that might otherwise be lost during conventional single-host enrichment workflows.

### 2.3. Direct spot test (DST)

Phage lytic activity was assessed using a DST. In this test, 100 μL of overnight bacterial cultures were added to 4 mL 0.5% (w/v) of soft agar, which was kept molten at 55 °C and then poured onto TSB plates. Subsequently, 10 μL of phage stock (10^9^ to 10^10^ PFU/mL) was spotted on the plates against a collection of 103 Staphylococcus strains, for subsequent DST-based host range analysis. The plates were incubated overnight at 37 °C. The results were categorised, scored as either a very clear lysis zone (Score 10), a clear lysis zone (Score 7.5) turbid lysis zone (Score 5), a very turbid/hazy lysis zone (Score 2.5), or no lysis zone (Score 0) [21, 22].

### 2.4. Efficiency of plating (EOP)-based host range analysis

From the panel of isolated phages, 28 representative phages were selected for host range and EOP analysis against a total of 103 *Staphylococcus* strains (70 *S. aureus* and 33 *S. epidermidis*). Overnight bacterial cultures were mixed with molten soft agar and overlaid onto TSA plates to generate bacterial lawns. For each strain, 5 µL aliquots of serially diluted phage suspensions (10^1^ to 10^8^ PFU/mL) were spotted onto the surface of the soft agar overlay. Plates were incubated at 37 °C for 24 h and examined for plaque formation. EOP values were determined by comparing the number of plaques formed on each test strain to those obtained on the corresponding reference host under similar conditions. EOP was calculated as the ratio of plaque-forming units (PFU) on the test strain to PFU on the reference strain. This approach was used to assess relative infectivity and host range across the full panel of *Staphylococcus* isolates. EOP values were classified as high (≥0.1), moderate (0.01-<0.1), low (0.001-<0.01), or no production (<0.001).

### 2.5. Phage adsorption assay and one-step growth assay

Phage adsorption kinetics and one-step growth curves were performed to determine adsorption efficiency, latent period, and burst size. Overnight bacterial cultures were diluted into fresh medium and grown to mid-log phase (OD_600_ ≈ 0.5; ∼1 × 10^8^ CFU/mL). Phages were added at low multiplicity of infection (MOI 0.01 or 0.1) to minimise multiple infections per cell.

For adsorption assays, phage-host mixtures were incubated at room temperature, and 200 μL aliquots were collected immediately (0 min) and after 5 and 10 min. Samples were centrifuged at 6000g for 1-2 min at 4 °C to pellet bacteria with adsorbed phages. Supernatants containing unadsorbed phages were removed, and pellets were washed once with SM buffer and resuspended to the original volume. At each time point, appropriate dilutions were prepared and plated using the double-layer agar method. Plaques were enumerated after incubation to estimate the proportion of adsorbed versus free phages, and dilution factors were adjusted to obtain countable plaques.

For one-step growth curve assays, bacterial cultures were infected at an MOI of 0.01 (or 0.1 for slow-adsorbing phages) and allowed to adsorb for a duration determined from prior adsorption kinetics experiments (1-3 mins). Following adsorption, cultures were centrifuged to remove unbound phages, and pellets were resuspended in fresh medium. The infected cultures were then diluted 1:10 to prevent secondary infections and incubated under the same growth conditions. At defined time points (0, 2, 4, 6, 8, 10, 15, 20, 30, 40, 50, and 60 min, 100 μL aliquots were withdrawn, serially diluted, mixed with soft agar and a 50 μL overnight bacterial host culture, and plated using the double-layer agar method. Plaques were counted after overnight incubation at 37°C to calculate the eclipse period and burst size.

### 2.6. Bacterial growth inhibition measured by OD_600_ nm kinetics

OD_600_-based killing assays were performed to evaluate phage-mediated growth inhibition and to optimise the MOI using a microplate reader (BMG LABTECH). Log-phase bacterial cultures were adjusted to a final concentration of ∼10^6^ CFU/mL, with a total assay volume of 200 µL per well. Phage preparations were added at varying MOIs to evaluate dose-dependent effects. Based on these preliminary experiments, an intermediate MOI (typically MOI=10) was selected, as it provided reproducible killing without saturation effects observed at very high MOIs or delayed inhibition at low MOIs. This optimised MOI was subsequently used for comparative killing assays.

### 2.7. Time-kill kinetics assay

Time-kill kinetics were evaluated using an initial bacterial inoculum of approximately 10^6^ CFU/mL. Briefly, 0.1 mL of a mid-exponential-phase bacterial culture (OD_600_= 0.5, corresponding to ∼10^8^ CFU/mL) was added to 9.9 mL of TSB. Phage suspensions were then added at a concentration sufficient to achieve an MOI of 10. Aliquots (200 µL) were collected immediately before phage addition (0 h) and at 2, 4, 6, and 24 h post-infection. Samples were centrifuged at 18,000 × g for 10 min at 4 °C, the supernatant was discarded, and bacterial pellets were resuspended in an equal volume of sterile normal saline (0.9% sodium chloride (NaCl) solution). Serial tenfold dilutions were prepared in 0.9%, and 50 µL of the appropriate dilutions were plated onto TSA. Viable bacterial counts were determined after incubation at 37 °C for 24 h. All time-kill experiments were performed independently for each phage-host combination.

### 2.8. Ion-supplemented serial passaging of phage-resistant strains

Highly phage-resistant *S. aureus* isolates identified from the host-range panel were selected for adaptive evolution experiments. Warrior phage (Φ21) and multiST-adapted phage K were incubated with resistant hosts at MOI = 1 in 1X TSB media containing 100 μL mid-log phase cultures supplemented with 10 mM MgSO_4_and 10 mM CaCl_2_. Cultures were incubated for 10 h, after which lysates were recovered, clarified, and used to infect fresh resistant cultures for subsequent passages. This serial passaging was repeated for five rounds. In parallel, control evolution experiments were performed under identical conditions without ionic supplementation. Evolved lysates were assessed by spot assay to evaluate changes in lysis clarity and halo formation.

### 2.9. Genomic analysis

Following the manufacturer’s instructions, phage genomic DNA was extracted using a commercial phage DNA isolation kit from Norgen Biotek. The quantity and purity of the DNA were measured using a NanoDrop spectrophotometer (Thermo Fisher Scientific). Whole-genome sequencing was performed on an Illumina HiSeq 2500 platform with paired-end 150 base pair (bp) reads, resulting in an average sequencing depth of approximately 100X (mean read coverage ranging from 95X to 105X). Raw sequencing reads were assembled *de novo* using SPAdes v4.0.0 software with default settings. Phage-specific functional annotation was further refined using the PHASTEST database. Comparative genomic analyses were conducted using the CaGeCaT (Comparative Genomics and Categorisation Tool; https://cagecat.bioinformatics.nl/) platform. Multiple sequence alignments for phylogenetic analysis were generated using the MAFFT software. All genome sequences have been uploaded to GenBank, and accession numbers are given in the Data Availability Statement.

### 2.10. Statistical analysis

All experiments were performed in biological triplicate unless otherwise stated. Raw data were recorded in Microsoft Excel and expressed as mean ± standard deviation. The relationship between eclipse period (min) and burst size (PFU per infected cell) was assessed using Pearson correlation analysis (two-tailed). Correlation strength was reported as Pearson r with 95% confidence intervals and coefficient of determination (R^2^). Differences in burst size and eclipse period between *S. aureus* and *S. epidermidis* phages were evaluated using exact two-tailed Mann–Whitney U tests due to non-normal distribution and unequal sample sizes. Medians are reported with Hodges–Lehmann estimates of median differences where appropriate. Statistical significance was defined as alpha = 0.05. Graphs and statistical analyses were conducted using GraphPad Prism (version 10.6.1) and Python (Google Colab). Figures and schematic visualisations were prepared using BioRender.

## 3. Results

A panel of fifty-six *Staphylococcus* isolates was initially subjected to phage isolation using conventional protocols, which are highly efficient for recovering phages targeting Gram-negative bacteria but are less effective for Gram-positive hosts [18]. This approach yielded only six unique phages, indicating limited recovery efficiency under standard conditions (**Supplementary File S1**). While conventional phage isolation protocols are highly effective for many Gram-negative hosts, recovery from Gram-positive bacteria can be reduced due to differences in cell surface architecture, including the absence of an outer membrane and the presence of a thick peptidoglycan layer that influences adsorption dynamics. This occurs due to variations in adsorption requirements that depend on the architecture of the cell envelope [2, 23, 24]. To address this challenge, we modified the phage isolation method for Gram-positive bacteria by providing a controlled ion environment, as detailed in **Section 2.2** and **Figure 1.1a**, to better support phage adsorption, early replication, and chloroform-assisted (or lysozyme-assisted for some challenging strains) release from the bacterial cells.

**Figure 1.**
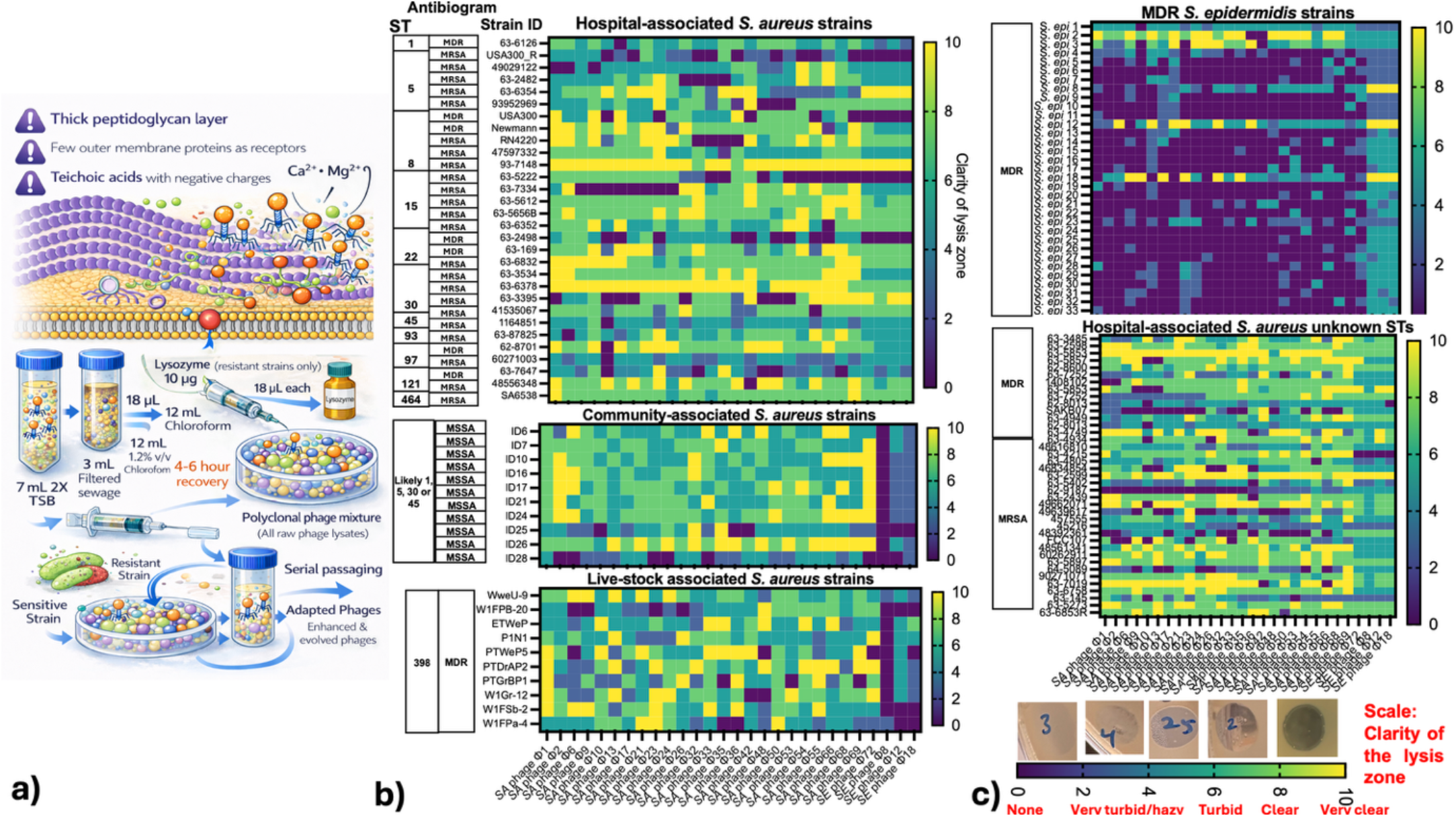
Optimised Gram-positive phage enrichment and host-range profiling. (a) Schematic of an enhanced phage isolation workflow incorporating Mg^2+^/Ca^2+^ supplementation, lysozyme treatment, and serial passaging to improve recovery of phages from MRSA and MDR clinical isolates, enabling development of a regional phage bank. (b) Heatmap of DST-based host-range profiling across major hospital-, community-, and livestock-associated *Staphylococcus aureus* sequence types, showing broad lytic activity against MSSA, MRSA, and MDR strains prevalent in Australia. (c) Additional host-range testing against MDR *Staphylococcus epidermidis* and hospital-associated *S. aureus* isolates demonstrates heterogeneous but widespread lytic activity across diverse resistance phenotypes.

By employing this method, we generated a panel of 28 genomically distinct novel lytic Staphylococcus phage isolates. Despite the reported complexity of phage susceptibility determinants in *S. epidermidis*, our optimised isolation framework enabled recovery of four lytic phages from a panel of 20 MDR isolates. Plaque morphologies ranged from small to large, with diameters between 1.2 and 7 mm, as detailed in **Supplementary File S1**. This substantial improvement in phage recovery confirms that ionic conditions critically shape the efficiency and success of phage isolation from Gram-positive hosts. The use of conventional phage isolation methods alone significantly underestimates the recoverable phage diversity. These findings show that ionic modulation and the use of polyclonal raw phage lysate mixtures are key determinants of phage recovery from Gram-positive bacteria. Divalent cation availability reshapes phage-host evolutionary dynamics and overcomes structural adsorption barriers in Gram-positive pathogens.

### 3.1. An ion-supplemented isolation strategy enabled broad sequence-type coverage by a collection of 28 novel lytic Staphylococcus phages

To establish a phage bank, we isolated 28 distinct Staphylococcal phages and evaluated their lytic activity across a diverse panel of clinical and livestock-associated bacterial isolates. The collection demonstrated broad activity against dominant Australian *S. aureus* sequence types (STs), including hospital-associated, community-associated, and livestock-associated (CC398) lineages (**Figure 1.2b**). Importantly, lytic activity was observed across both MRSA and multidrug-resistant (MDR) strains, supporting the translational potential of the collection as a therapeutic resource for clinically challenging infections. The phage panel was also tested against a batch of MDR *S. epidermidis* isolates with very challenging and complex antibiotic susceptibility profiles. Despite challenging antibiograms, several *S. epidermidis* strains showed susceptibility to multiple phages. Together, these results indicate that the 28-phage collection provides broad functional coverage across Staphylococcal lineages circulating in both Australian healthcare and livestock settings.

### 3.2. High EOP phage clusters: ion-supported adsorption & productive infection

Combined analysis of spot-test and EOP data across 103 *Staphylococcus* strains (70 *S. aureus*, 33 *S. epidermidis*) revealed interesting lytic trends, indicating a heterogeneous infection landscape determined by each phage-host interaction. (**Supplementary Figure S1a**). Across all 2,884 phage-host combinations, spot-test screening showed a broad spectrum of lytic outcomes, with 15.0% very clear lysis, 26.5% clear lysis, 20.3% turbid lysis, 11.8% very turbid/hazy lysis, and 26.4% showing no detectable lysis, indicating that ∼41.5% of interactions produced well-defined lysis zones, either clear or turbid. These qualitative trends were later validated by quantitative EOP measurements, where overall 51.7% of interactions showed high infectivity (EOP ≥ 0.5), 16.6% medium production (0.01-<0.1), 1.1% low production (0.001-<0.01), and 30.6% no or inefficient production (<0.001), confirming that plaque clarity broadly reflects underlying infection efficiency. Significant species-level differences were also observed. *S. aureus* supported efficient phage replication in the majority of interactions, with 68.6% high-EOP outcomes, 16.1% medium (0.01-<0.1), and only 15.4% non-productive interactions. In contrast, *S. epidermidis* showed resistance towards phage treatments, exhibiting (165/247; 60%) non-productive, medium, or low-productive EOP outcomes. These trends closely resemble spot-test results, where ∼62% of *S. epidermidis* interactions showed no visible lysis, compared with ∼10-15% for *S. aureus*, indicating strong host-dependent barriers to productive infection.

### 3.3. One-step growth curves reveal heterogeneity in replication kinetics

One-step growth experiments demonstrated significant heterogeneity in replication dynamics across the Staphylococcal phages (**Figure 2**). Eclipse periods ranged from 20 to 40 min, and burst sizes varied from 3 to 72.1 PFU per infected cell (**Figure 2a-c, Supplementary figure S1b;**). A subset of phages combined short eclipse periods (20-25 min) with high progeny production, including Φ35 (72.1 PFU/infected cell), multiST-adapted phage K (65 PFU per infected cell), Warrior phage Φ21 (64 PFU per infected cell), Φ32 (62 PFU per infected cell), and *S. epidermidis* phage Φ72 (64 PFU per infected cell), consistent with rapid intracellular replication and efficient host lysis. In contrast, several phages exhibited intermediate burst sizes (approximately 20-40 PFU per infected cell), while others showed limited phage copies each burst (<15 PFU per infected cell) despite comparable eclipse periods, indicating reduced replication efficiency (**Figure 2a-c**). Across the dataset, burst size was significantly inversely correlated with eclipse period (Pearson r = −0.443, p = 0.016), indicating that phages with shorter eclipse periods generally produced higher progeny yields (**Figure 2d**). However, no statistically significant differences in burst size or eclipse period were observed between phages infecting different *Staphylococcus* species (*S. aureus*, n = 25; *S. epidermidis*, n = 4) (p = 0.3485 and p = 0.5929, respectively), suggesting that replication kinetics were primarily phage-specific rather than determined by the host *Staphylococcus* species (**Figure 2e**).

**Figure 2.**
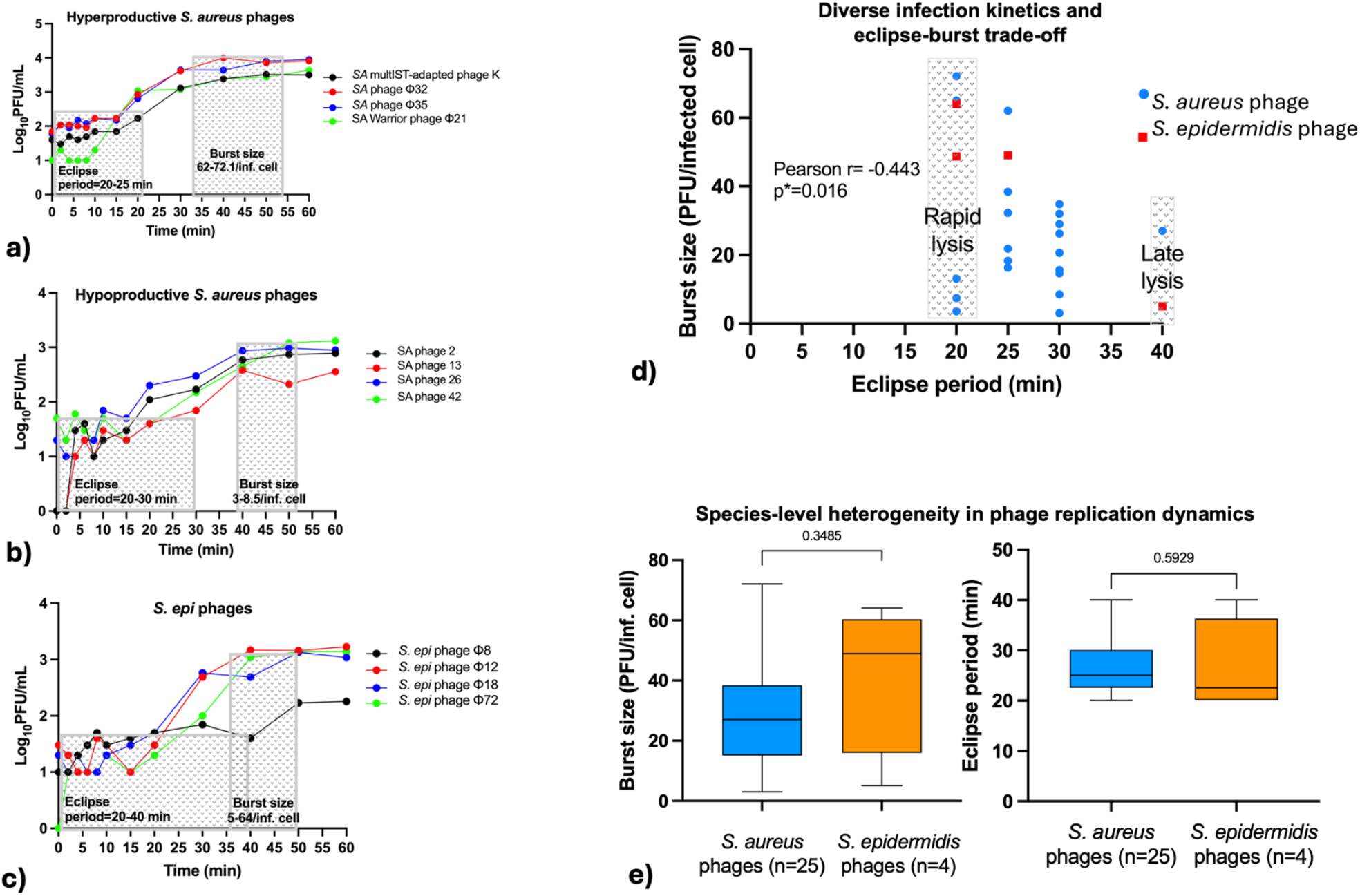
Replication kinetics of Staphylococcus phages. (**a-c**) One-step growth curves show diverse replication phenotypes, with hyperproductive *S. aureus* phages displaying short eclipse periods and large burst sizes, while hypoproductive phages exhibit lower amplification. *S. epidermidis* phages show variable burst sizes. (**d**) Burst size was inversely correlated with eclipse period (r = -0.443, p = 0.016), indicating a replication trade-off. (**e**) No significant differences in burst size or eclipse period were observed between *S. aureus* and *S. epidermidis* phages, suggesting replication dynamics are largely phage-intrinsic.

### 3.4. The majority of Staphylococcus phages, specifically “Warrior Phage” showed a cross-species broad host range

Isolated phage host ranges were quantified as the proportion of strains supporting productive infection (EOP ≥ 0.5) across the 103-isolate panel (**Figure 3a**). A significant heterogeneity in infectivity and host range was observed among the phages. The broadest host range was exhibited by *S. aureus* phage Φ21 or the Warrior phage, which productively infected 63.1% of strains. This was followed by Φ68 (58.3%), Φ54 (56.3%), Φ69 and Φ50 (each 55.3%), and Φ55 (53.4%). Several additional phages also demonstrated relatively broad host ranges, including Φ9 and Φ24 (both 53.4%), Φ42 (52.4%), Φ53 and Φ66 (51.4%), and Φ23 (52.4%). A large group of phages displayed intermediate infectivity, with host ranges around 48-50%, including Φ1, Φ6, Φ17, Φ32, Φ33, and Φ35 (48.54-49.5%). More restricted host ranges were observed for Φ10 and Φ26 (44.7%), Φ36 (35%), and Φ2 (42.7%).

**Figure 3.**
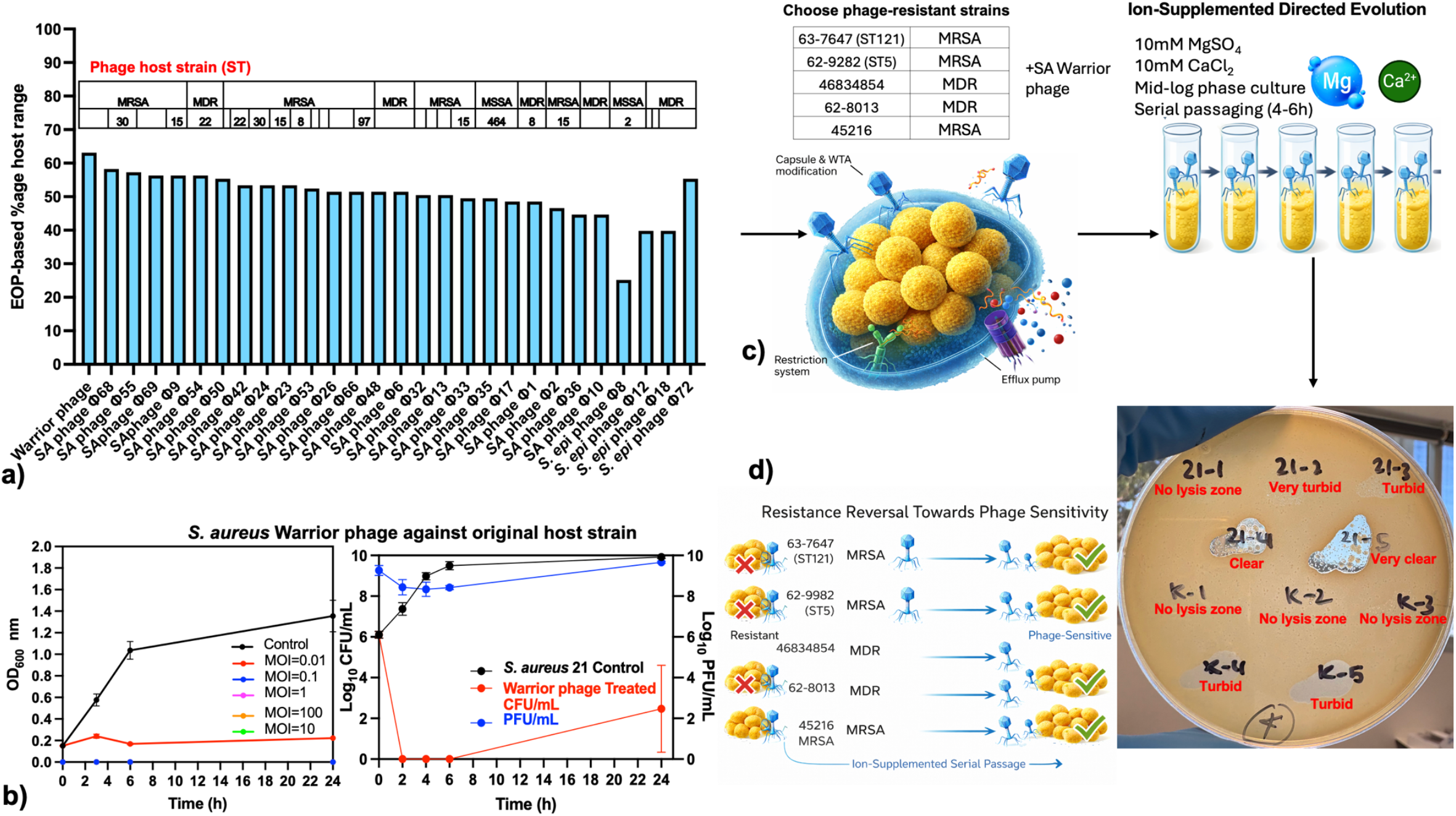
Ion-supplemented directed evolution expands phage host range. (**a**) Host-range ranking of phages based on EOP-defined productive infection across MRSA and MDR *Staphylococcus* sequence types. (**b**) Time-kill analysis of the Warrior phage shows dose-dependent bacterial suppression over 24 h. (**c-d**) Ion-supplemented serial passaging with Mg^2+^/Ca^2+^ restores infectivity against previously resistant strains. (e) Spot assays confirm improved lytic activity following ion-directed evolution.

Among *S. epidermidis* phages, Φ72 exhibited a broad host range (55.3%), comparable to several of the most infective *S. aureus* phages. In contrast, Φ8 showed a narrow host range (25.2%) because the majority (n=26/103; 77%) of phage Φ8-bacterial infection displayed either low infectivity (0.001-<0.01) and no or inefficient infectivity (45%), while Φ12 and Φ18 displayed moderate host ranges of ∼40%. Overall, our data highlight significant variability in host-range breadth across the phage panel, with half of the total phages (16/28; 57.1%) exhibiting broad-spectrum (>50% coverage) activity across diverse *Staphylococcus* isolates. Most *S. epidermidis* isolates exhibited substantial resistance to both *S. aureus* and *S. epidermidis* phages, clearly demonstrated by small and turbid/hazy plaque formation. In contrast, the majority of *S. aureus* isolates remained broadly susceptible to the Staphylococcal phage panel (**Supplementary figure S1a)**.

### 3.5. Phage-mediated killing is largely independent of MOI

Across the 28 phages tested, interesting trends were observed in killing kinetics, host range, and dose dependence. **Figure 3b** shows that the warrior phage treatment resulted in rapid and sustained suppression of bacterial growth at all MOIs (0.01 to 100) compared with the untreated control, with significant reductions in viable CFU within the first few hours post-infection. PFU titres increased over time, indicating productive phage replication within the host population throughout 24h course of the infection cycle. Likewise, most phages showed minimal effects at early time points but induced more noticeable growth suppression by 24 h, consistent with sustained killing activity (**Supplementary Figure S1c**).

Higher MOIs generally showed rapid early killing; however, sustained suppression at later time points was frequently maintained even at lower EOP, indicating efficient phage amplification. Interestingly, some phages exhibited MOI-independent killing at 24 h, with comparable suppression across a wide range of doses. These included *S. aureus* phages Φ1, Φ6, Φ9, Φ10, Φ13, Φ17, Warrior phage Φ21, Φ23, Φ24, Φ26, Φ32, Φ33, Φ35, Φ36, Φ42, Φ48, Φ50, Φ53, Φ54, Φ55, Φ66, Φ68, and Φ69, which consistently reduced bacterial growth even at MOI ≤ 1, suggesting high adsorption efficiency and strong killing kinetics and phage amplification. In contrast, other phages displayed partial killing, indicated by regrowth at 24 h and stronger dependence on higher MOIs. Phages infecting *S. epidermidis* exhibited greater variability overall, with delayed killing kinetics and stronger MOI dependence. OD_600_-based time-kill curves at MOI=10 supported these dose-independent killing trends, showing rapid suppression within 3-6 h and sustained inhibition at 24 h for highly active phages, while revealing partial regrowth for phages with low EOP. Collectively, our findings highlight a broad landscape of infection phenotypes, ranging from highly potent phages, demonstrated by sustained killing activity and no growth 24h post-phage treatment, to phages with intermediate or low killing, shown by regrowth at either later or early timepoints, respectively.

### 3.6. Ion-supplemented directed evolution restores and enhances lytic activity against phage-resistant hosts

This ion-supplemented directed evolution approach enables expansion of host range and restoration of lytic activity without the need to isolate new phages, providing a rapid and scalable strategy to adapt existing phages against resistant hosts. To overcome resistance barriers, we subjected four highly phage-resistant *S. aureus* strains to serial passaging with Warrior phage (Φ21) and phage multiST-adapted K (**Figure 3c)**. Ion-supplemented evolution (10 mM Mg^2+^ and Ca^2+^) resulted in progressive improvement in lysis clarity over five sequential passages, with visibly expanded and more defined halo zones compared to the parental phages (**Figure 3c and d**). All four resistant strains exhibited enhanced susceptibility following ion-directed evolution. In contrast, evolution performed without ionic supplementation also produced modest improvements in lysis, but halo clarity and expansion were consistently reduced relative to the ion-supplemented condition. The upper panel of the plate (ion-supplemented evolution) in **Figure 3d** demonstrates a more visible clearing or lysis zone compared to the lower panel (conventional (non-ionic evolution), with each spot representing a successive passage. These findings indicate that divalent cation-mediated serial passaging accelerates adaptive recovery of infectivity in resistant Staphylococcal hosts, consistent with the established role of Ca^2+^ and Mg^2+^ in adsorption efficiency and phage-host binding dynamics.

### 3.7. South Australia’s first phage bank: A regional platform for phage treatments against clinical and livestock-associated Staphylococcus species

Our findings establish South Australia’s first phage bank as a regional or local platform with the capacity to provide broad therapeutic coverage across the wide array of *S. aureus* STs (ST1, ST5, ST8, ST15, ST22, ST30, ST45, ST93, ST97, ST121, ST464) (**Figures 1b and 4a-d**) prevalent in the Australian population as well as most dominant livestock-associated lineage CC398 (**Figure 4e**). In collaboration with South Australian healthcare facilities, we established a continuously expanding phage bank supported by a continuous supply of MDR and extensively drug-resistant (XDR) clinical isolates, enabling future development of therapeutic-graded monophage treatment options and phage cocktails aligned with regional epidemiology. Quantitative time-kill kinetics integrating OD_600_, CFU/mL, and PFU/mL measurements demonstrated sustained bactericidal activity across dominant Australian sequence types. Multiple novel lytic phages effectively suppressed HA-MRSA ST15 and ST22 and CA-MRSA ST30 (**Fig. 4b-d**). Likewise, the Warrior phage exhibited potent activity against LA-MRSA CC398 isolates of porcine origin (**Fig. 4e**), underscoring cross-sectoral relevance across human and livestock settings.

**Figure 4.**
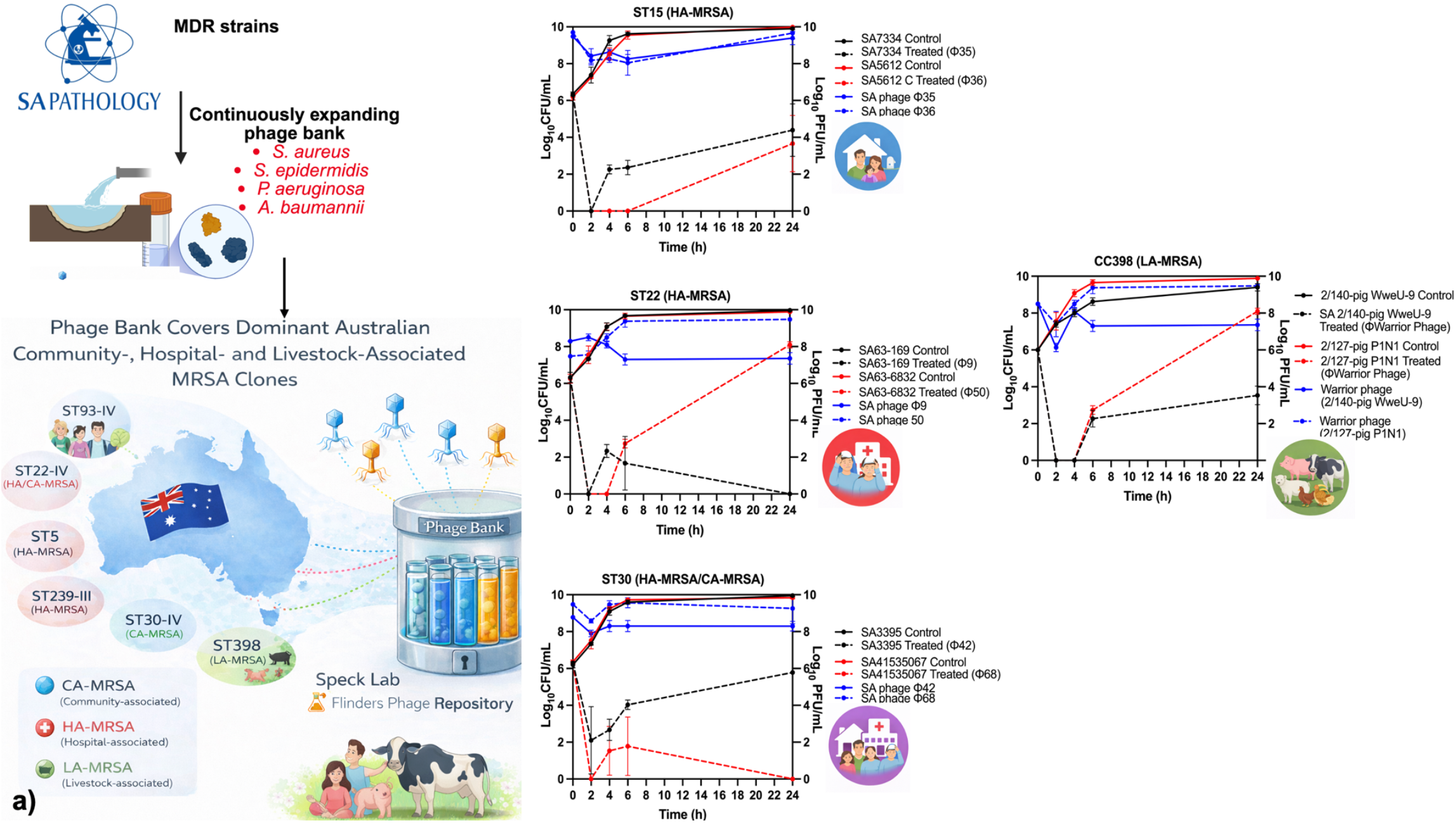
Regional phage bank shows broad therapeutic activity against major *S. aureus* lineages. (**a**) Overview of the South Australian phage repository covering key hospital-, community-, and livestock-associated *S. aureus* sequence types. (**b-d**) Time-kill assays demonstrate effective bacterial suppression and phage amplification against clinically relevant MRSA lineages. (**e**) The Warrior phage also shows strong activity against livestock-associated MRSA CC398 isolates, supporting cross-host efficacy.

### 3.8. Phage-mediated bacterial killing dynamics indicate sustained lytic activity

To validate and quantitatively refine the growth and killing patterns observed in OD_600_-based assays, we next performed time-kill (TK) experiments measuring growth inhibition as CFU/mL and phage replication dynamics as PFU/mL at an MOI of 10 (**Supplementary figure S1d**). Unlike OD measurements, TK assays provide a more sensitive and accurate direct assessment of bactericidal activity, allowing discrimination between true killing, delayed growth, and regrowth driven by resistant subpopulations. The results of the TK dataset analysis confirmed that many phage treatments showed sustained killing activity, demonstrated by significant reductions in bacterial burden compared to untreated controls. In particular, phages Φ1, Φ6, Φ9, Φ10, Φ13, Φ21, Φ23, Φ24, Φ26, Φ32, Φ36, Φ48, Φ54, Φ66, Φ68, and Φ69 consistently showed no growth relative to controls at 24h timepoint. Likewise, several phages such as Φ2, Φ17, Φ21, Φ33, Φ42, Φ50, Φ53, Φ8, Φ12, Φ18, and Φ72 exhibited partial regrowth during the 4-6 h post-infection window, yet bacterial densities remained significantly reduced compared with the untreated control at 24 h.

### 3.9. Phage replication dynamics demonstrate sustained persistence

Time-dependent PFU measurements showed that all phages remained detectable throughout the 24 h infection period, indicating sustained persistence during bacterial challenge (**Supplementary figure S1f**). Several phages maintained high titres approaching ∼10^9^ PFU/mL at 24 h post-treatment, reflecting efficient replication, persistence, and stability over time. Transient reductions in PFU/mL at early time points (2-6h) were observed for some phages such as Φ9, Φ13, Φ17, Φ48, and Φ55. Although the phage persistence profile and magnitude of amplification varied among phages and host backgrounds, no phage exhibited rapid loss of infectivity during the assay. Overall, phage persistence and continuous efficient replication are supported by the sustained high PFU levels observed 24h post-phage treatment, which also reflect a mechanistic explanation for the sustained bacterial growth inhibition observed in the corresponding CFU time-kill tests.

### 3.10. Genomic architecture shows that host-range variation correlates with divergence in tail-associated genes

Comparative genomic analysis of our *Staphylococcus* phage collection showed a conserved Kayvirus backbone typical of obligately lytic Herelleviridae. Genome size (133-140 kb), coding capacity (193-216 CDS), GC content (∼30%), and replication-lysing modules are highly conserved across the panel, indicating functional similarities in DNA replication, morphogenesis, and host lysis across *S. aureus* and *S. epidermidis* phages (**Table 1** and **Figure 5a**).

**Table 1.** Genomic characterisation of the novel lytic *Staphylococcus* phages isolated in this study.

| Phage ID | Genome size (bp) | No of CDS | GC content (%) | Structural proteins | Non-structural proteins | Best blast hit (% identity coverage) |
| --- | --- | --- | --- | --- | --- | --- |
| **S. aureus phage Φ1** | 140,025 | 216 | 30.40 | Capsid and Head proteins, Tail associated proteins, Membrane proteins | Endolysins, Endonucleases, Helicase, Polymerases, Reductases, Peptidoglycan binding proteins, transcriptase, holin, terminase | 99.62%<br>Staphylococcus phage 812<br>Herelleviridae, Kayvirus |
| **S. aureus phage Φ2** | 133,709 | 198 | 30.45 | Capsid and Head proteins, Tail associated proteins, Membrane proteins | Endolysins, Endonucleases, Helicase, Polymerases, Reductases, Peptidoglycan binding proteins, transcriptase, holin, terminase | 99.36%<br>Staphylococcus phage Staph1N<br>Herelleviridae, Kayvirus |
| **S. aureus phage Φ6** | 134,086 | 199 | 30.44 | Capsid and Head proteins, Tail associated proteins, Membrane proteins | Endolysins, Endonucleases, Helicase, Polymerases, Reductases, Peptidoglycan binding proteins, transcriptase, holin, terminase | 99.24%<br>Staphylococcus phage Staph1N<br>Herelleviridae, Kayvirus |
| **S. aureus phage Φ9** | 133,919 | 200 | 30.45 | Capsid and Head proteins, Tail associated proteins, Membrane proteins | Endolysins, Endonucleases, Helicase, Polymerases, Reductases, Peptidoglycan binding proteins, transcriptase, holin, terminase | 99.36%<br>Staphylococcus phage Staph1N<br>Herelleviridae, Kayvirus |
| **S. aureus phage Φ10** | 137,987 | 193 | 29.90 | Capsid and Head proteins, Tail associated proteins, Membrane proteins | Endolysins, Endonucleases, Helicase, Polymerases, Reductases, Peptidoglycan binding proteins, transcriptase, holin, terminase | 97.1%<br>Staphylococcus phage St1.CM<br>Unclassified |
| **S. aureus phage Φ13** | 133,795 | 199 | 30.45 | Capsid and Head proteins, Tail associated proteins, Membrane proteins | Endolysins, Endonucleases, Helicase, Polymerases, Reductases, Peptidoglycan binding proteins, transcriptase, holin, terminase | 99.24%<br>Staphylococcus phage Staph1N<br>Herelleviridae, Kayvirus |
| **S. aureus phage Φ17** | 140,340 | 216 | 30.37 | Capsid and Head proteins, Tail associated proteins, Membrane proteins | Endolysins, Endonucleases, Helicase, Polymerases, Reductases, Peptidoglycan binding proteins, transcriptase, holin, terminase | 99.37%<br>Staphylococcus phage 812<br>Herelleviridae, Kayvirus |
| **S. aureus phage Φ21** | 134,247 | 200 | 30.45 | Capsid and Head proteins, Tail associated proteins, Membrane proteins | Endolysins, Endonucleases, Helicase, Polymerases, Reductases, Peptidoglycan binding proteins, transcriptase, holin, terminase | 99.65%<br>Staphylococcus phage SAML-229<br>Unclassified |
| **S. aureus phage Φ23** | 137,985 | 193 | 29.9 | Capsid and Head proteins, Tail associated proteins, Membrane proteins | Endolysins, Endonucleases, Helicase, Polymerases, Reductases, Peptidoglycan binding proteins, transcriptase, holin, terminase | 97.18%<br>Staphylococcus phage St1.CM<br>Unclassified |
| **S. aureus phage Φ24** | 133,004 | 197 | 30.47 | Capsid and Head proteins, Tail associated proteins, Membrane proteins | Endolysins, Endonucleases, Helicase, Polymerases, Reductases, Peptidoglycan binding proteins, transcriptase, holin, terminase | 99.48%<br>Staphylococcus phage Staph1N<br>Herelleviridae, Kayvirus |
| **S.**<br>**aureus**<br>**phage**<br>**Φ26** | 139,649 | 214 | 30.40 | Capsid and<br>Head proteins,<br>Tail associated<br>proteins,<br>Membrane<br>proteins | Endolysins,<br>Endonucleases, Helicase,<br>Polymerases, Reductases,<br>Peptidoglycan binding<br>proteins, transcriptase,<br>holin, terminase | 99.20%<br>Staphylococcus<br>phage 812<br>Herelleviridae,<br>Kayvirus |
| **S.**<br>**aureus**<br>**phage**<br>**Φ32** | 133,513 | 196 | 30.41 | Capsid and<br>Head proteins,<br>Tail associated<br>proteins,<br>Membrane<br>proteins | Endolysins,<br>Endonucleases, Helicase,<br>Polymerases, Reductases,<br>Peptidoglycan binding<br>proteins, transcriptase,<br>holin, terminase | 98.55%<br>Staphylococcus<br>phage phiSNIID<br>Herelleviridae,<br>Kayvirus |
| **S.**<br>**aureus**<br>**phage**<br>**Φ33** | 134,296 | 199 | 30.99 | Capsid and<br>Head proteins,<br>Tail associated<br>proteins,<br>Membrane<br>proteins | Endolysins,<br>Endonucleases, Helicase,<br>Polymerases, Reductases,<br>Peptidoglycan binding<br>proteins, transcriptase,<br>holin, terminase | 99.24%<br>Staphylococcus<br>phage Staph1N<br>Herelleviridae,<br>Kayvirus |
| **S.**<br>**aureus**<br>**phage**<br>**Φ35** | 134,355 | 199 | 30.43 | Capsid and<br>Head proteins,<br>Tail associated<br>proteins,<br>Membrane<br>proteins | Endolysins,<br>Endonucleases, Helicase,<br>Polymerases, Reductases,<br>Peptidoglycan binding<br>proteins, transcriptase,<br>holin, terminase | 99.35%<br>Staphylococcus<br>phage Staph1N<br>Herelleviridae,<br>Kayvirus |
| **S.**<br>**aureus**<br>**phage**<br>**Φ36** | 139,053 | 214 | 30.42 | Capsid and<br>Head proteins,<br>Tail associated<br>proteins,<br>Membrane<br>proteins | Endolysins,<br>Endonucleases, Helicase,<br>Polymerases, Reductases,<br>Peptidoglycan binding<br>proteins, transcriptase,<br>holin, terminase | 99.32%<br>Staphylococcus<br>phage 812<br>Herelleviridae,<br>Kayvirus |
| **S.**<br>**aureus**<br>**phage**<br>**Φ42** | 138,354 | 193 | 29.91 | Capsid and<br>Head proteins,<br>Tail associated<br>proteins,<br>Membrane<br>proteins | Endolysins,<br>Endonucleases, Helicase,<br>Polymerases, Reductases,<br>Peptidoglycan binding<br>proteins, transcriptase,<br>holin, terminase | 97.18%<br>Staphylococcus<br>phage St1.CM<br>Unclassified |
| **S.**<br>**aureus**<br>**phage**<br>**Φ48** | 140,598 | 216 | 30.40 | Capsid and<br>Head proteins,<br>Tail associated<br>proteins,<br>Membrane<br>proteins | Endolysins,<br>Endonucleases, Helicase,<br>Polymerases, Reductases,<br>Peptidoglycan binding<br>proteins, transcriptase,<br>holin, terminase | 99.34%<br>Staphylococcus<br>phage 812<br>Herelleviridae,<br>Kayvirus |
| **S.**<br>**aureus**<br>**phage**<br>**Φ50** | 140,313 | 216 | 30.39 | Capsid and<br>Head proteins,<br>Tail associated<br>proteins,<br>Membrane<br>proteins | Endolysins,<br>Endonucleases, Helicase,<br>Polymerases, Reductases,<br>Peptidoglycan binding<br>proteins, transcriptase,<br>holin, terminase | 99.18%<br>Staphylococcus<br>phage 812<br>Herelleviridae,<br>Kayvirus |
| **S.**<br>**aureus**<br>**phage**<br>**Φ53** | 140,313 | 215 | 30.41 | Capsid and<br>Head proteins,<br>Tail associated<br>proteins,<br>Membrane<br>proteins | Endolysins,<br>Endonucleases, Helicase,<br>Polymerases, Reductases,<br>Peptidoglycan binding<br>proteins, transcriptase,<br>holin, terminase | 99.26%<br>Staphylococcus<br>phage 812<br>Herelleviridae,<br>Kayvirus |
| **S.**<br>**aureus**<br>**phage**<br>**Φ54** | 140,143 | 197 | 30.44 | Capsid and<br>Head proteins,<br>Tail associated<br>proteins,<br>Membrane<br>proteins | Endolysins,<br>Endonucleases, Helicase,<br>Polymerases, Reductases,<br>Peptidoglycan binding<br>proteins, transcriptase,<br>holin, terminase | 98%<br>Staphylococcus<br>phage Staph1N<br>Herelleviridae,<br>Kayvirus |
| **S.**<br>**aureus**<br>**phage**<br>**Φ55** | 133,810 | 197 | 30.46 | Capsid and<br>Head proteins,<br>Tail associated<br>proteins,<br>Membrane<br>proteins | Endolysins,<br>Endonucleases, Helicase,<br>Polymerases, Reductases,<br>Peptidoglycan binding<br>proteins, transcriptase,<br>holin, terminase | 97.18%<br>Staphylococcus<br>phage St1.CM<br>Unclassified |
| **S.**<br>**aureus**<br>**phage**<br>**Φ66** | 138,319 | 198 | 30.44 | Capsid and<br>Head proteins,<br>Tail associated<br>proteins,<br>Membrane<br>proteins | Endolysins,<br>Endonucleases, Helicase,<br>Polymerases, Reductases,<br>Peptidoglycan binding<br>proteins, transcriptase,<br>holin, terminase | 99.19%<br>Staphylococcus<br>phage 812<br>Herelleviridae,<br>Kayvirus |
| **S.**<br>**aureus**<br>**phage**<br>**Φ68** | 134,016 | 193 | 29.90 | Capsid and<br>Head proteins,<br>Tail associated<br>proteins,<br>Membrane<br>proteins | Endolysins,<br>Endonucleases, Helicase,<br>Polymerases, Reductases,<br>Peptidoglycan binding<br>proteins, transcriptase,<br>holin, terminase | 99.39%<br>Staphylococcus<br>phage Staph1N<br>Herelleviridae,<br>Kayvirus |
| **S.**<br>**aureus**<br>**phage**<br>**Φ69** | 137,985 | 197 | 30.46 | Capsid and<br>Head proteins,<br>Tail associated<br>proteins,<br>Membrane<br>proteins | Endolysins,<br>Endonucleases, Helicase,<br>Polymerases, Reductases,<br>Peptidoglycan binding<br>proteins, transcriptase,<br>holin, terminase | 97.18%<br>Staphylococcus<br>phage St1.CM<br>Unclassified |
| **S.**<br>**aureus**<br>**phage**<br>**Φ72** | 133,953 | 193 | 29.90 | Capsid and<br>Head proteins,<br>Tail associated<br>proteins,<br>Membrane<br>proteins | Endolysins, Endonuclease,<br>Helicase, Polymerases,<br>Reductases,<br>Peptidoglycan binding<br>proteins, transcriptase,<br>holin, terminase | 99.67%<br>Staphylococcus<br>phage SAML-229<br>Unclassified |

**Figure 5.**
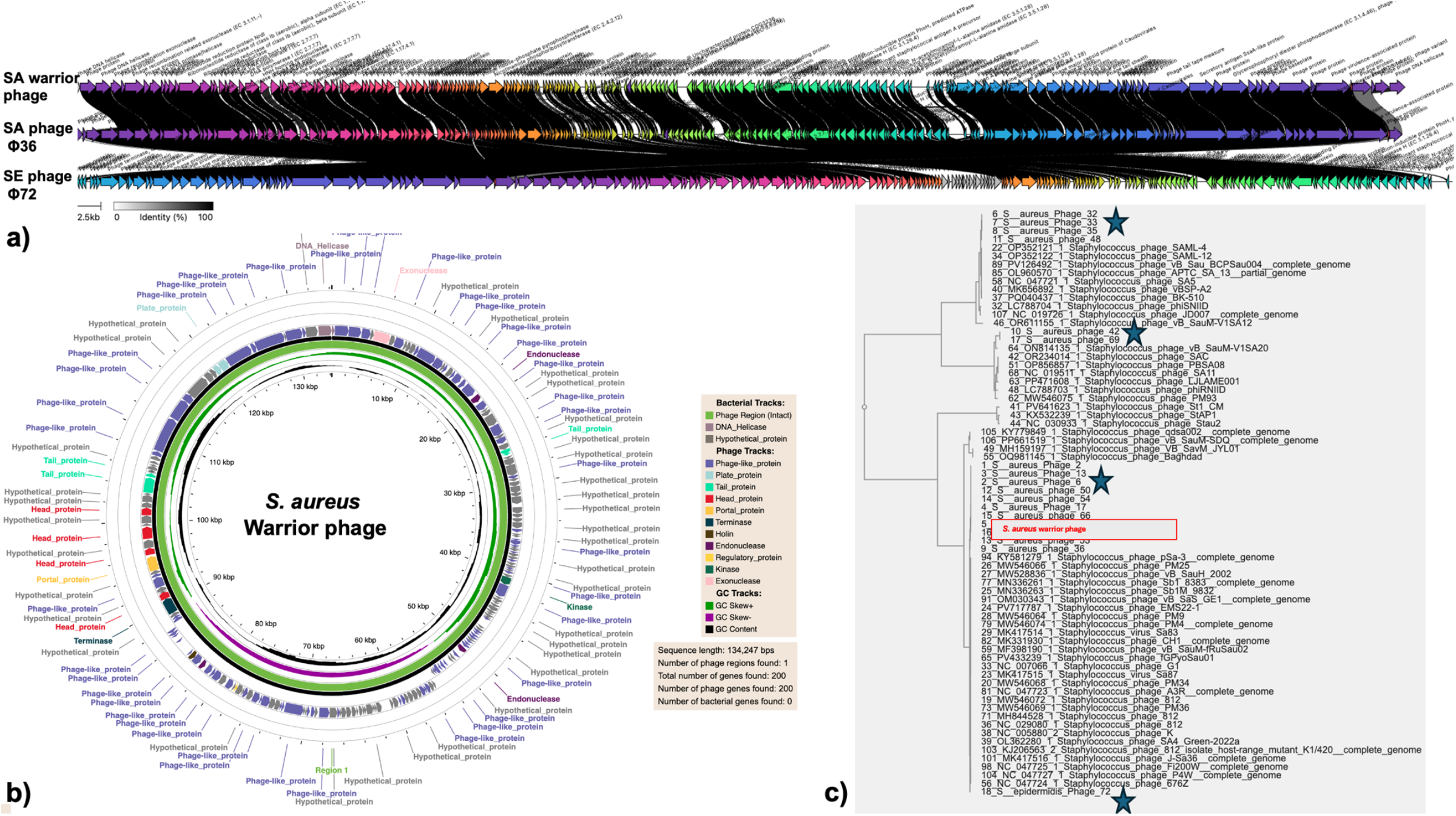
Regional phage bank shows broad therapeutic activity against major *S. aureus* lineages. (**a**) Overview of the South Australian phage repository covering key hospital-, community-, and livestock-associated *S. aureus* sequence types. (**b-d**) Time-kill assays demonstrate effective bacterial suppression and phage amplification against clinically relevant MRSA lineages. (**e**) The Warrior phage also shows strong activity against livestock-associated MRSA CC398 isolates, supporting cross-host efficacy.

Within our panel, Warrior phage Φ21 demonstrated the strongest killing efficiency and the broadest host range (**Figure 4b**). The circular genomic annotation of all phages are given in **Supplementary Figure S2**. Representative genomic annotation (**Supplementary File S2; Figure 5b**) shows that Warrior phage Φ21 encodes dual holins, two endolysins, a CHAP-domain protein, N-acetylmuramoyl-L-alanine amidase, and LysM-domain peptidoglycan-binding proteins. Although LysM-containing proteins are present across the phage collection, differences in gene regulation and expression may contribute to the enhanced bactericidal activity observed for Warrior phage Φ21.

The presence of multiple, functionally distinct lytic enzymes likely increases the efficiency of cell wall degradation, which may underlie its superior killing phenotype. Another phage Φ36, produced the largest plaques among all phage isolates in the panel. SA phage Φ36 encodes ribonucleotide reductase (NrdE/NrdI) together with an expanded replication module comprising multiple DNA polymerases, primase, helicase loader, and DNA repair proteins, suggesting increased nucleotide autonomy and potentially faster intracellular replication. This genomic configuration likely underlies its bigger plaque morphology through enhanced replication. However, despite forming large plaques, Φ36 displayed only moderate killing efficiency and a comparatively narrower host range. Together, these findings suggest that plaque size alone does not predict the phage’s killing potential.

Phylogenetic analysis showed that our phage panel was classified into four major clusters within the Kayvirus group (**Figure 5c**). Cluster I comprised Φ32, Φ33, Φ35, and Φ48, which grouped closest to *Staphylococcus phage* SAML-4, indicating strong similarity to this reference lineage. A second cluster containing Φ42 and Φ69 is positioned between reference genomes *vb_SauM-V1SA20* and *vb_SauM-V1SA12*. Cluster III included Φ2, Φ13, Φ6, Φ50, Φ54, Φ17, Φ16, and Φ36, was found grouped between *Staphylococcus phage Baghdad* and *Staphylococcus phage pSa-3*, indicating shared ancestry within this genomic framework. Warrior phage Φ21 (warrior phage) resides within this broader lineage. In contrast, the *S. epidermidis* phage clustered separately at the base of the tree, closest to *Staphylococcus phage 676Z*, reflecting deeper divergence from the *S. aureus* lineages.

## 4. Discussion

Bacteriophages are increasingly recognised as promising biological agents shaping microbial diversity and combating AMR, yet phage host interactions in Gram-positive bacteria remain mechanistically underexplored. In Staphylococcus species, infection is primarily constrained at the adsorption and genome delivery stages due to the thick, highly cross-linked peptidoglycan layer and structurally diverse wall teichoic acids that function as principal phage receptors. Variations in teichoic acid backbone composition, glycosylation, and surface charge directly modulate receptor accessibility and binding affinity, making phage discovery and recovery inherently more challenging than in Gram-negative systems. Although our Kayvirus panel shared conserved replication and lysis modules, including multiple holins, endolysins, CHAP domain proteins, and LysM peptidoglycan binding elements, productive infection was ultimately governed by efficient adsorption and DNA injection through this restrictive cell wall architecture (**Figure 5, Supplementary File S2**). We demonstrate that many limitations attributed to biological resistance are in fact methodological barriers imposed by conventional isolation protocols [18]. By optimising ionic conditions, enrichment using polyclonal raw lysates, and host physiological states, we established a collection of 28 lytic Staphylococcus phages (**Figure 1a; Supplementary File S1**). Divalent cations such as Ca^2+^ and Mg^2+^ mechanistically stabilise tail fibre interactions with negatively charged teichoic acids, reduce electrostatic repulsion, and facilitate DNA translocation, markedly improving phage recovery. For a subset of highly resistant strains, low-concentration lysozyme was applied during the initial enrichment step to mildly loosen the peptidoglycan layer without disrupting receptor architecture [25, 26]. Furthermore, ion-supplemented serial passaging accelerated adaptive restoration and expansion of lytic activity against resistant hosts, providing a rapid and scalable strategy to enhance infectivity and broaden host range without the need for time-consuming de novo phage isolation. This mechanistically informed framework supports the development of an expanding therapeutic-grade phage repository in collaboration with SA Pathology.

Direct evidence for the ionic supplementation of media as a standardised strategy for bacteriophage isolation remains limited; however, several studies have demonstrated that ionic composition plays a critical role in phage adsorption, genome injection, and early infection dynamics [27, 28]. Likewise, in another study, phages isolated from sewage and applied as mixtures have been shown to exhibit enhanced activity when supplemented with divalent metal ions; for example, Ca^2+^ supplementation improved phage adsorption, productivity, and antibacterial efficacy of a *Staphylococcus aureus* phage cocktail both *in vitro* and *in vivo*, without altering host range [29]. This effect is particularly relevant for Gram-positive bacteria, including *Staphylococcus* spp., where the thick peptidoglycan layer and abundant teichoic acids present substantial physicochemical barriers to phage attachment and entry [14, 30]. Divalent cations such as Ca^2+^ and Mg^2+^ are known to facilitate phage-host interactions by shielding negatively charged cell surface polymers, stabilising adsorption complexes, and promoting conformational rearrangements in phage tail and baseplate structures required for productive infection [31, 32]. Our phage isolation strategy, optimised for Gram-positive phages, also confirmed that prolonged enrichment incubations can impose strong selective pressure favouring phage-resistant bacterial subpopulations or rapidly replicating phage variants with narrow host specificity, potentially biasing recovery outcomes (**Figure 1**). Consistent with this, our workflow employed short enrichment periods (≤6 h), sufficient to allow productive infection and phage amplification while minimising overgrowth of resistant hosts that commonly emerge during overnight incubations. Finally, the use of a polyclonal phage reservoir derived from pooled raw lysates preserves low-abundance and genetically diverse phage populations, enabling repeated host-specific enrichment without repeated environmental sampling. Together, the integration of mid-log-phase hosts, optimised ionic conditions, limited incubation time, and a shared phage reservoir constitutes a rational, selection-aware strategy that enhances recovery efficiency and supports sustained expansion of a diverse *Staphylococcus* phage library.

The combined DST and EOP analyses demonstrate that qualitative lysis does not always reflect efficient phage replication (**Figure 1b and 3a; Supplementary File S2**). While spot tests are useful for rapid screening, visible clearing can overestimate functional host range, as some interactions showing lysis did not translate into strong phage production. This highlights the importance of incorporating quantitative EOP assays when defining therapeutic host range [33, 34]. The species-level differences discussed in **Figure 2d and e** further suggest strong host-dependent barriers to productive infection. Phages replicated more efficiently in *S. aureus*, whereas *S. epidermidis* frequently exhibited restricted or inefficient production, likely due to differences in surface receptors or intracellular defence mechanisms (**Supplementary File S1**). Together, these findings emphasise that robust host range characterisation requires combined qualitative and quantitative assessment to avoid overestimating phage efficacy [35, 36]. These observations are consistent with previous comparative host range studies showing that spot assays often overestimate virulence and breadth when not supported by EOP measurements. Similar discrepancies between qualitative lysis and productive infection have been reported in ECOR-based analyses, where single dilution spot tests failed to correlate with true replication efficiency. Similar strain-dependent reductions in EOP have been seen for other Myophages, including phage GE1 and phage K, where high apparent susceptibility in spot assays did not translate into consistent productive infection and EOP values [34, 37]. Our data extend this concept to broad range of Staphylococci strains, reinforcing the need for strict EOP-based validation when selecting candidates for therapeutic phage libraries [38].

Our comparatively low recovery of *S. epidermidis* phages, with only four isolates displaying narrow host ranges, likely reflects the intrinsic biological resistance of the strain panel rather than a limitation of the isolation strategy (**Figure 1b and 3a**). As highlighted by Beck et al. [17], *S. epidermidis* susceptibility is tightly governed by wall teichoic acid architecture, glycosylation state, and a diverse repertoire of phage defence systems, including restriction modification and abortive infection mechanisms [39, 40]. The structured and receptor-specific nature of *S. epidermidis* phage interactions inherently restricts broad replication across strains. In this context, successful isolation of even a limited number of lytic phages from a highly refractory clinical batch underscores the stringency of our screening framework and highlights the robustness of our workflow against a genetically and phenotypically resistant *S. epidermidis* population.

Our findings position ion-supplemented directed evolution as a major translational strength of this study (**Figure 3c and d**). Laboratory-based phage training strategies such as the Appelmans protocol have been used to expand host range, including in carbapenem resistant *Acinetobacter baumannii*, but these approaches often depend on prolonged adaptation cycles, recombination events, or prophage mobilisation, and can yield derivatives with limited stability [41]. Likewise, hospital-specific optimisation strategies, such as pre-adaptation of phages against *Enterobacter cloacae* complex to increase isolate coverage, require repeated passaging and targeted isolation over extended timeframes to achieve broader efficacy [42]. Similarly, adaptive evolution of phages against biofilm-embedded *Pseudomonas aeruginosa* has shown that improved infectivity frequently arises from mutations in tail fibre and baseplate genes that enhance adsorption to heterogeneous surface receptors [43]. Collectively, these studies identify adsorption as the principal bottleneck limiting phage efficacy. In contrast to these multi-week systems, our ion-directed evolution framework achieved marked restoration and enhancement of lytic activity within five sequential passages over only five days, even against highly phage-resistant Staphylococcal strains [44-46]. By supplementing Ca^2+^ and Mg^2+^ during serial passaging, we mechanistically enhanced adsorption efficiency and stabilised phage host binding dynamics, thereby increasing the probability of selecting beneficial receptor interaction variants under controlled conditions [45, 46]. Importantly, we coupled this with early phage recovery at each passage rather than prolonged overnight amplification, thereby minimising overgrowth of resistant bacterial subpopulations and maintaining selective pressure toward improved infectivity rather than resistance dominance [46]. The visibly expanded and clarified lysis zones observed under ion-supplemented conditions, compared with modest improvements under non-ionic evolution, demonstrate that divalent cations actively accelerate adaptive recovery of infectivity [47]. Together, this strategy enables rapid expansion of effective host range and recovery of lytic potency without further phage isolation, offering a scalable and clinically relevant approach for personalised treatment against refractory clinical isolates without any delays that may sometimes cause death of critically ill patients [48, 49].

This study provides the first evidence of a local phage bank in South Australia with broad-spectrum phages targeting *S. aureus* and *S. epidermidis* sequence types prevalent across Australian community, hospital, and livestock settings (**Figure 4**). Importantly, several community-associated phages in our collection demonstrated strong lytic activity against livestock-associated *S. aureus*, including LA-MRSA CC398 pig isolates. LA-MRSA CC398 is now recognised as established and widespread in Australian piggeries, where it represents a significant reservoir of multidrug-resistant staphylococci [50]. Consistent with this epidemiological concern, the CC398 strains within our panel exhibited extensive multidrug resistance profiles and carried multiple AMR genes [51]. The ability of our phages to effectively suppress these highly resistant livestock-associated isolates highlights their potential utility in settings where antibiotic treatment options are limited. The impact extends beyond individual infections. Given the documented capacity of CC398 to move between animal and human populations, effective phage activity against these MDR livestock strains supports a broader One Health strategy aimed at reducing antimicrobial use in agriculture, limiting amplification of resistance within animal reservoirs, and mitigating zoonotic spillover into community and hospital environments.

### Translational impact and clinical application of this phage platform in future

A major strength of this study is the establishment of South Australia’s first continuously expanding Staphylococcus phage bank, developed in collaboration with infectious disease experts of SA Pathology. Direct and continuous access to MDR clinical isolates enables routine phage discovery and real-time surveillance of commonly circulating clones. By integrating systematic phage isolation, comprehensive host-range profiling, quantitative time-kill kinetics, and adaptive evolutionary optimisation, we generated broad-spectrum phage collections and rationally designed cocktails providing approximately 95% coverage across Australian *S. aureus* STs and MDR *S. epidermidis* strains. This transforms phage discovery from a benchside research activity into a structured, regionally aligned translational framework. Through this study, we are also highlighting the need to develop a proper infrastructure to produce therapeutic-grade phages and bridge the gap between benchside phage discovery to bedside clinical translational pipeline.

The clinical relevance of this platform is highlighted by the ongoing burden of invasive MRSA infections in Australia, particularly bacteraemia and sepsis, where delays in effective therapy are associated with significant morbidity and mortality [48, 49]. At present, patients in South Australia requiring compassionate phage therapy depend on interstate programs, specifically the phage bank in New South Wales, for phage treatment options. However, phage susceptibility is highly strain-specific, and in some cases, no suitable match has been identified for locally circulating isolates. In one recent South Australian case involving an MDR infection, interstate screening did not yield an active phage, and the patient died because treatment could not be delivered in time. These observations emphasise the importance of a local clinical phage bank. By combining continuous clinical isolate access, broad baseline coverage, and the use of optimised methods to deal with challenging bacterial strains, this study lays the foundation for a South Australia–based clinical translational pipeline capable of delivering strain-specific phage therapies within timeframes compatible with rapidly progressing infections. With continued expansion and alignment to regional epidemiology, this framework has the potential to develop into a nationally coordinated precision phage system for severe MDR infections, specifically Staphylococcus infections.

## Conclusion

In summary, this study demonstrates that many of the challenges traditionally associated with isolating and characterising phages targeting Gram-positive bacteria arise from methodological constraints rather than intrinsic biological limitations. By optimising ionic conditions, host physiological state, enrichment duration, and the use of polyclonal phage reservoirs, we established a robust and reproducible framework for recovering diverse, highly lytic Staphylococcus phages active against MSSA, MRSA, MDR *S. aureus*, and MDR *S. epidermidis*. Broad host range, cross-lineage infectivity, and sustained killing activity were consistently achieved when adsorption dynamics and adaptive pressures were strategically controlled. Importantly, the demonstrated activity of community-derived phages against livestock-associated MDR lineages, including LA-MRSA CC398, extends the relevance of this work beyond hospital settings. LA-MRSA CC398 is established within Australian piggeries and represents both a veterinary health challenge and a zoonotic concern. Given the economic significance of pig production to Australia’s livestock and food systems, effective control of MDR *S. aureus* in livestock has implications not only for antimicrobial stewardship but also for production sustainability and food security. By addressing resistant strains circulating across healthcare, livestock, and food production environments, this study supports a One Health approach that recognises the interconnected nature of human health, animal health, and agricultural economics.

## Acknowledgement

The project was funded by NHMRC Ideas Grant 2036550. We thank Professor Howard Fallowfield and SA Water for providing sewage samples used in this study. We also acknowledge SA Pathology for supplying bacterial strains. Sequencing services were provided by BMKGENE (Beijing, China) through their multi-omics sequencing platform. We acknowledge members of the Speck Laboratory (Dr. Felise Adams, Dr. Maoge Zang (Lawrence), Ms. April ven der Kamp and Ms. Anjali Bandara for sharing bacterial subculture stocks. The manuscript underwent language editing with the assistance of Grammarly Premium.

## Authors’ contribution

B.W. conceived and designed the study, performed the experimental work, analysed the data, and wrote the manuscript. J.Z. conducted all bioinformatic and genomic analyses. A.T. and M.M. critically reviewed the study design and provided methodological guidance. J.R. provided technical support relevant to the supply of bacterial strains and assisted with experimental work. M.W. and P.S. supervised the project, contributed to the study design and reviewed and edited the manuscript. All authors discussed the results and approved the final version of the manuscript.

The authors declare no conflict of interest.

## Data availability statement

All data generated or analysed during this study are included in this published article and its supplementary information files. Genome sequence data have been deposited in the NCBI GenBank database under the accession numbers BioProject PRJNA1425730 (will be released after publication).

