## Supplementary material for "Ionic regulation of Gram-positive phage adsorption governs host range and improves phage isolation efficiency": SupplementaryFigure1.docx

**a)**
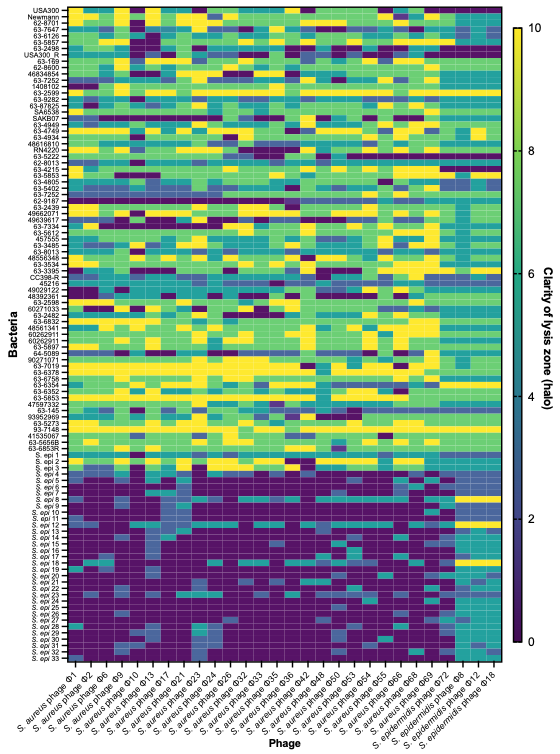

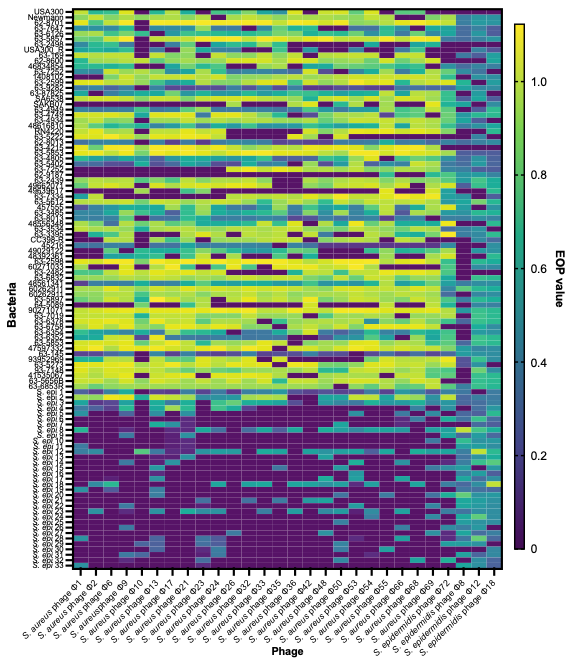

**b)**
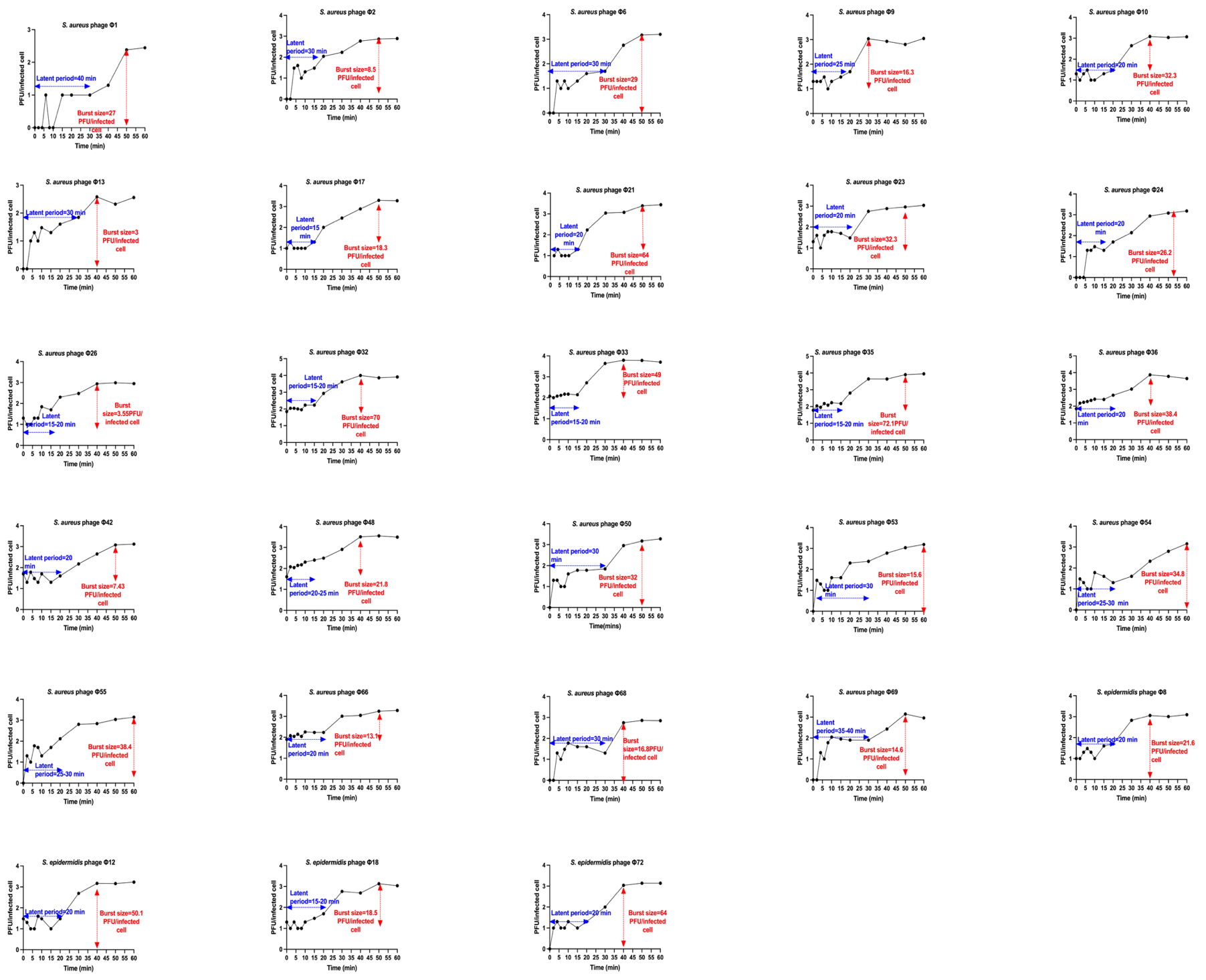

**c)**
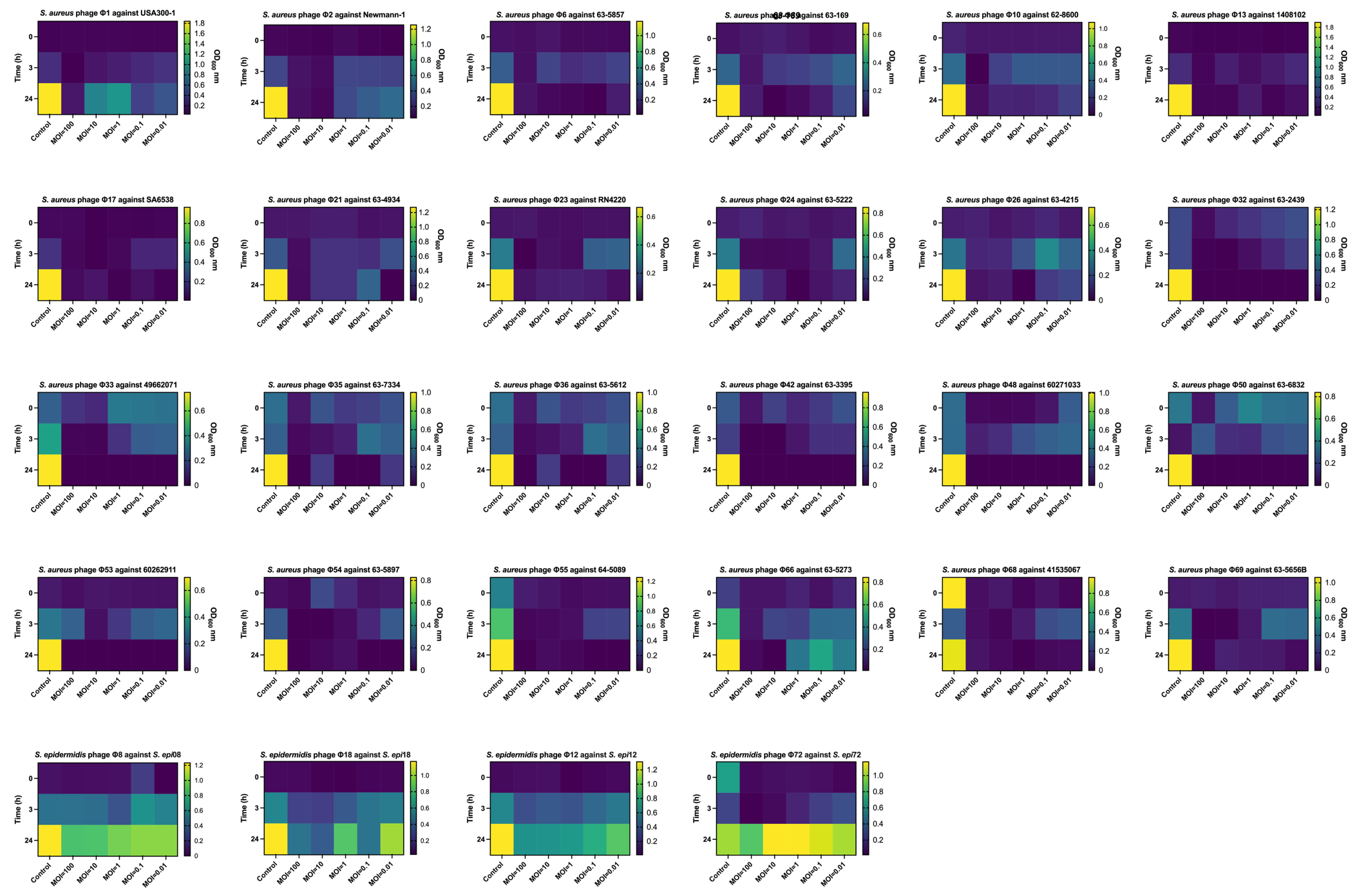

d)
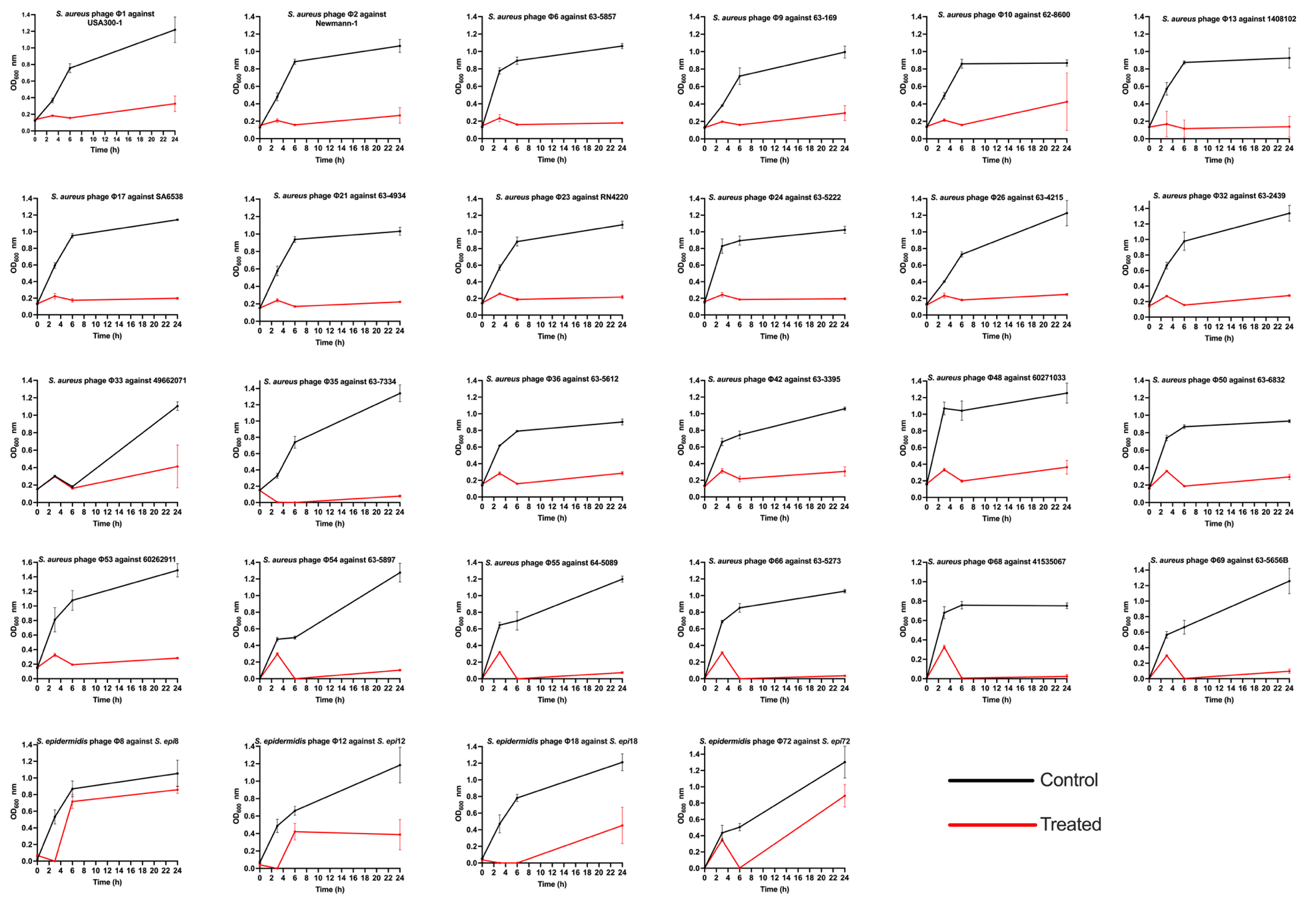

e)
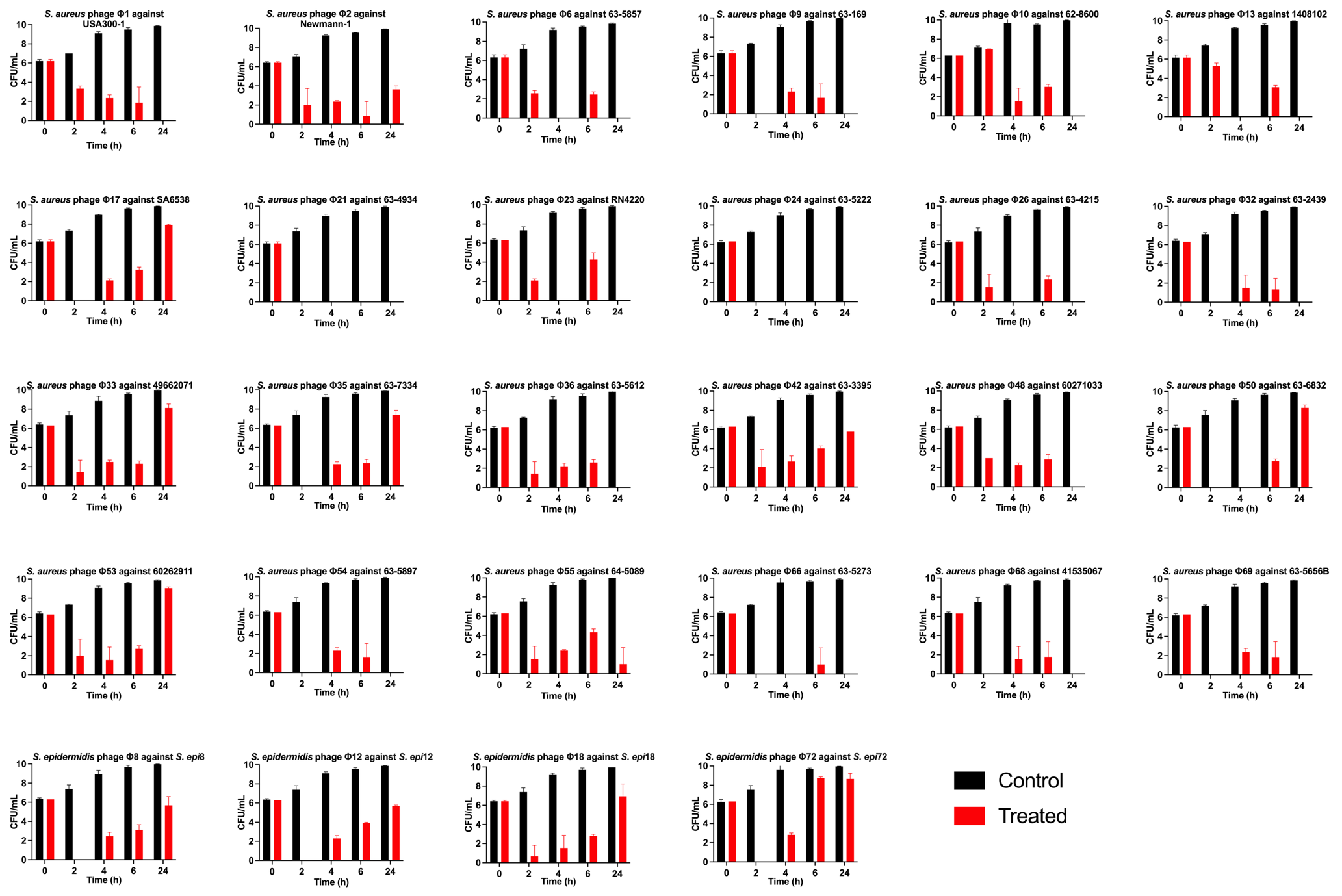

f)
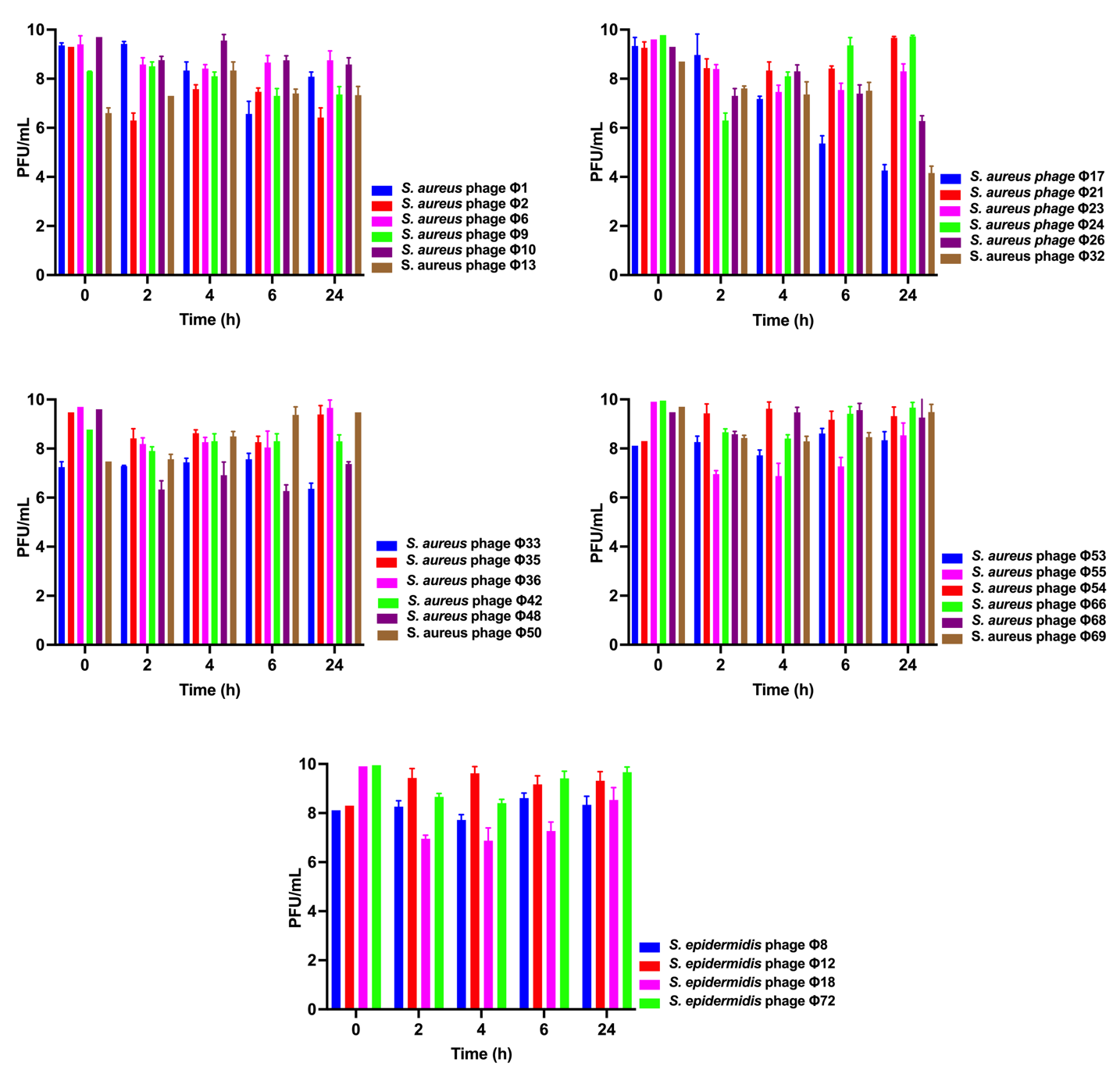

g)
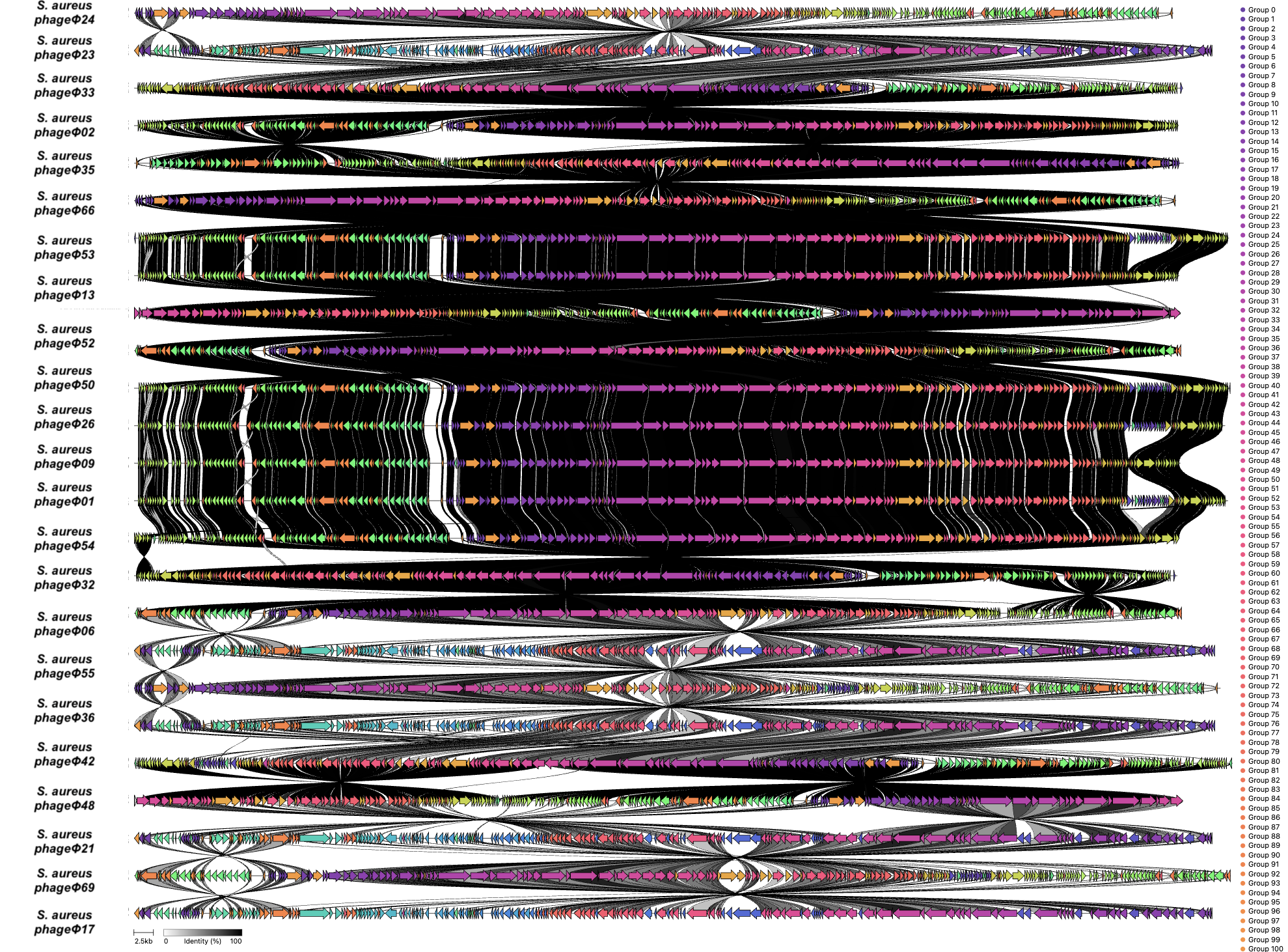

**Supplementary Figure 1: Comprehensive phenotypic and genomic characterisation of Staphylococcal bacteriophages.**

**a)** Analysis of lytic capacity of novel lytic phages against a panel of 103 MSSA, MRSA, and MDR Staphylococcus strains. Direct-spot test-based screening and EOP-based screening of the same isolate panel. **b)** Representative one-step growth curves demonstrating eclipse period and burst size for individual S. aureus and S. epidermidis phages. Blue arrows indicate the estimated latent period, while red arrows denote calculated burst size (PFU per infected cell). Latent periods ranged approximately 15–40 min, with burst sizes between 3–72 PFU per infected cell. **c)** Time- and MOI-dependent killing dynamics. Heatmaps show bacterial growth (OD_600_ nm) over time following infection with individual phages at MOIs of 0.01, 0.1, 1, 10, and 100. Time (0–24 h) is displayed on the y-axis and MOI on the x-axis. Colour intensity reflects residual bacterial growth, where lower OD indicates stronger killing. Untreated controls show uninhibited growth. **d)** OD600-based killing kinetics at MOI 10. Growth curves show the effect of individual phages on S. aureus and S. epidermidis strains. Experiments were performed in triplicate and presented as mean ± SD. **e)** Time–kill kinetics assessing bacterial viability (CFU/mL) over 24 h following phage treatment compared with untreated controls. Several phages induced rapid multi-log reductions and sustained suppression. Data represent mean ± SD from triplicate experiments. **f)** Phage replication dynamics (PFU/mL) during 24 h infection. Phage titres remained elevated, with several isolates reaching or maintaining approximately 10^9 PFU/mL at later time points, indicating productive replication. Data represent mean ± SD from triplicate experiments. **g)** Comparative genome alignment of broad-host-range Staphylococcus bacteriophages. Whole-genome synteny comparison of representative novel lytic S. aureus phages generated using pairwise nucleotide similarity mapping. Horizontal tracks represent complete genomes with ORFs coloured by functional category. Black ribbons denote regions of nucleotide similarity, with darker shading indicating higher identity.
