## Supplementary material for "Ionic regulation of Gram-positive phage adsorption governs host range and improves phage isolation efficiency": SupplementaryFigure2.docx

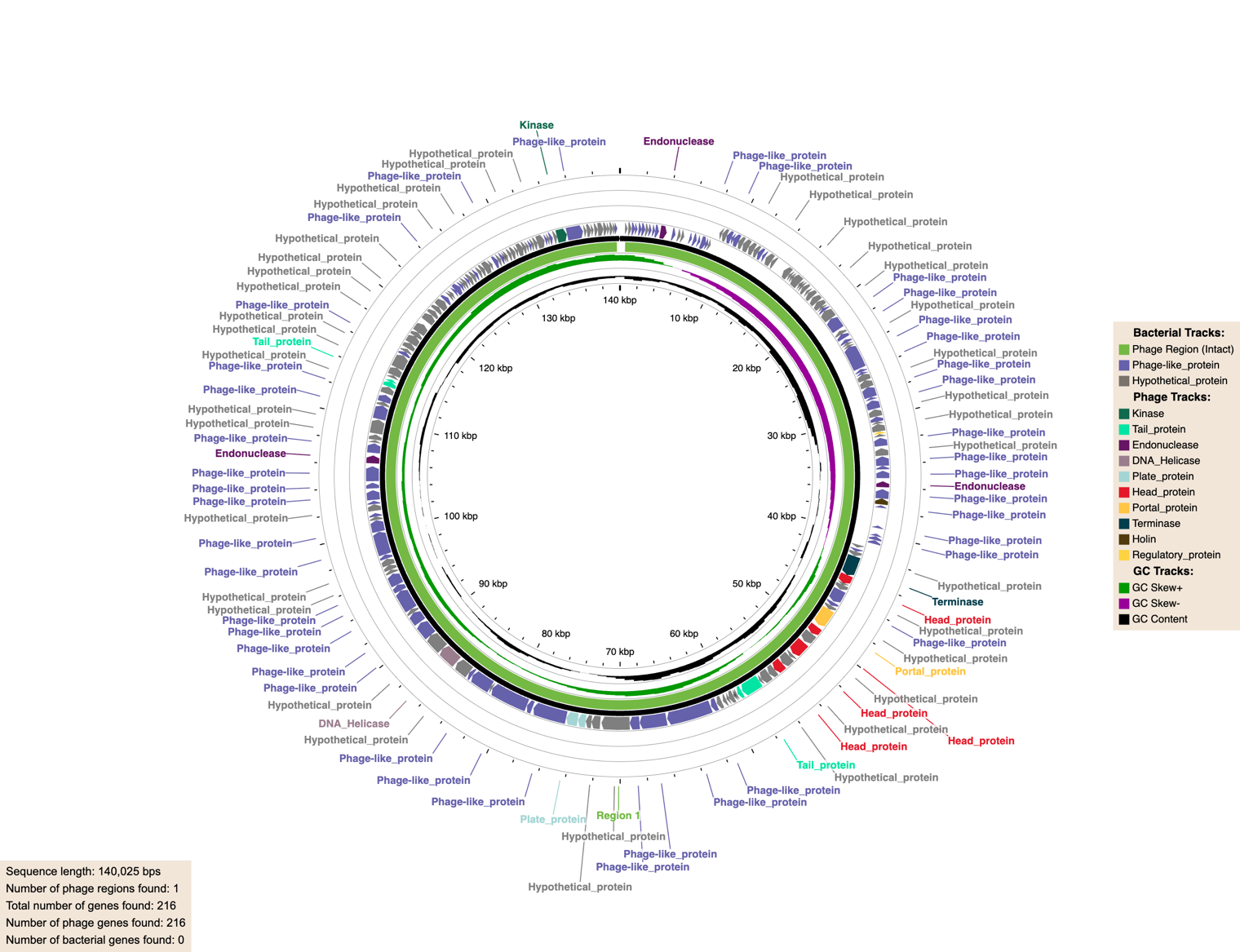

**a)**

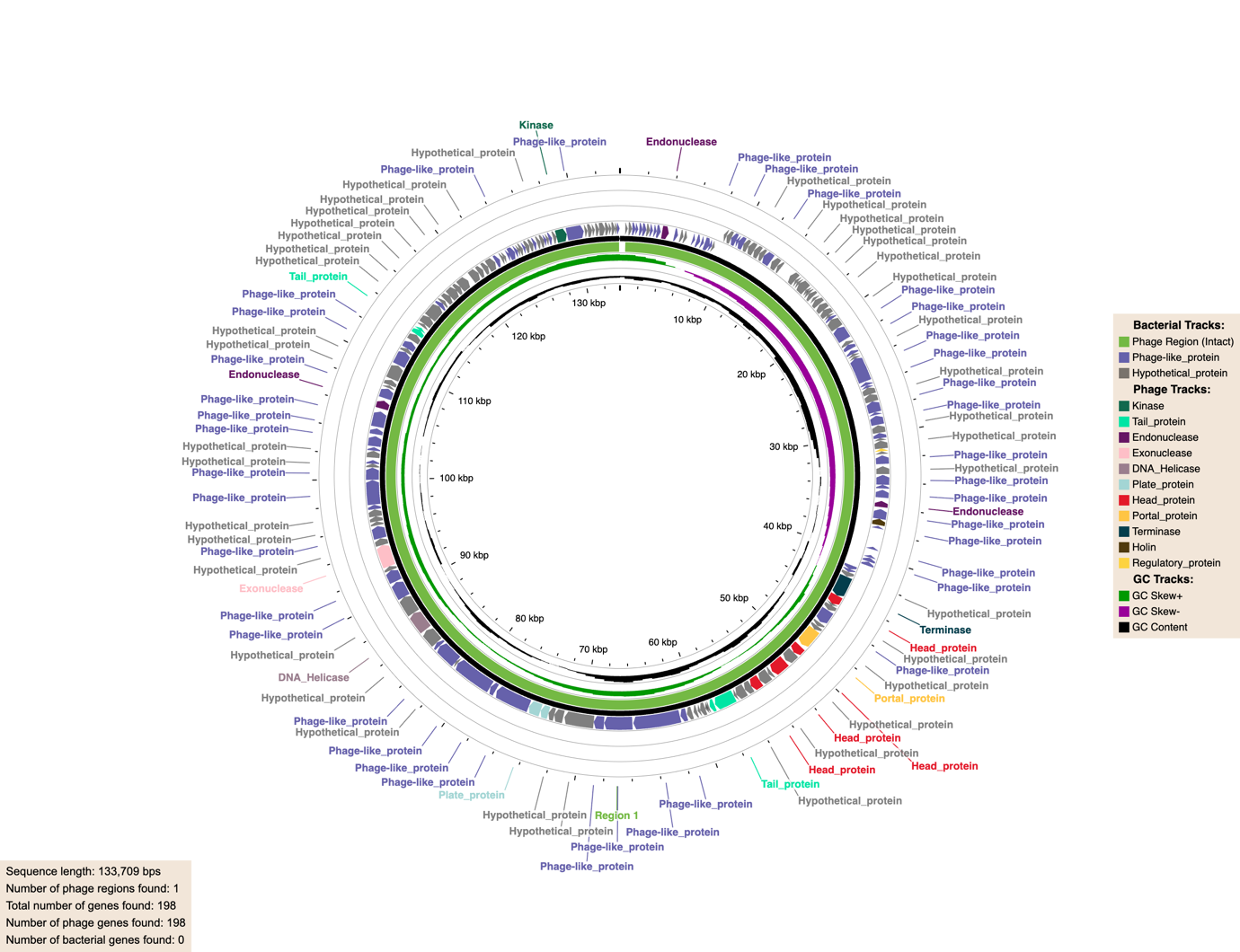

**b)**

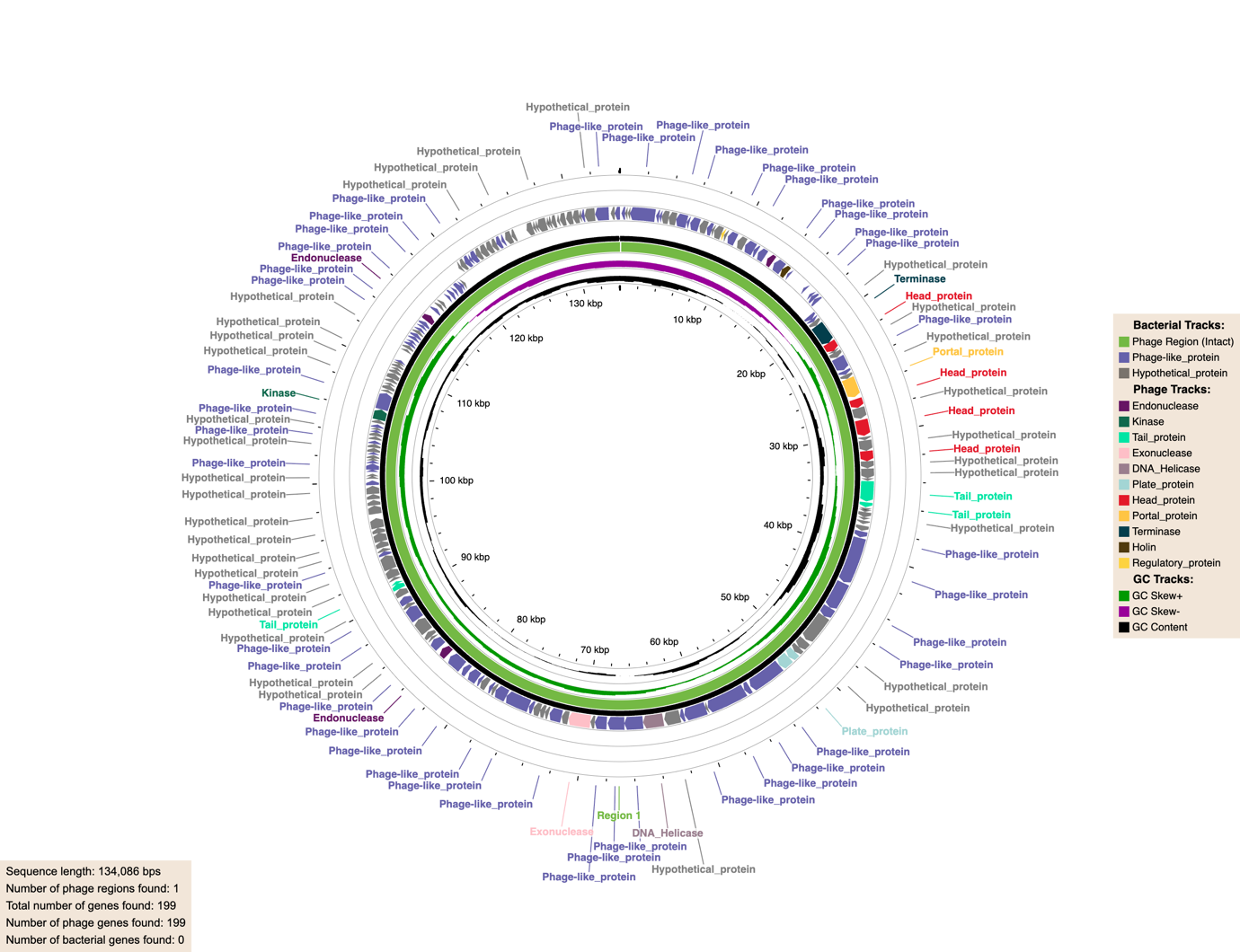

**c)**

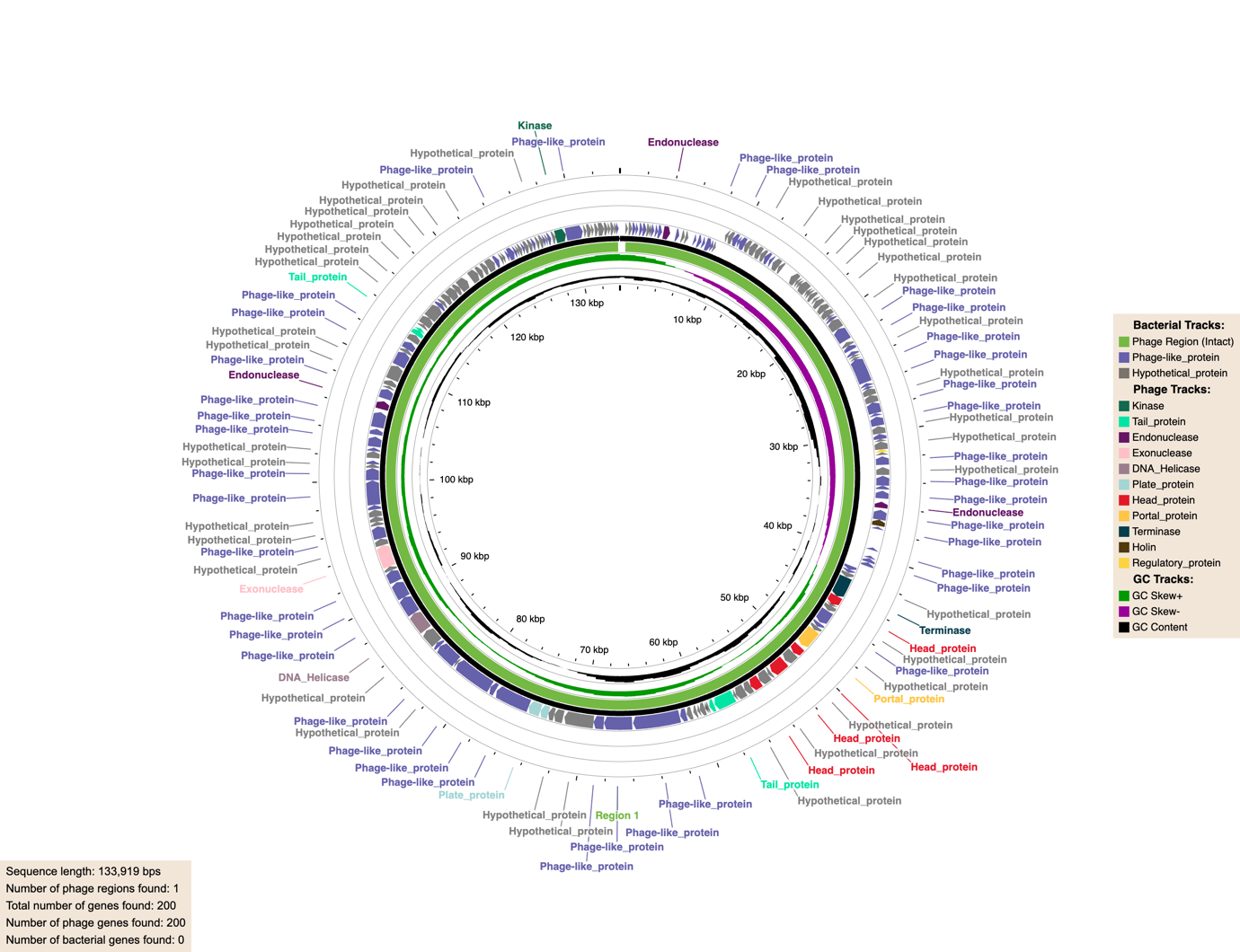

**d)**

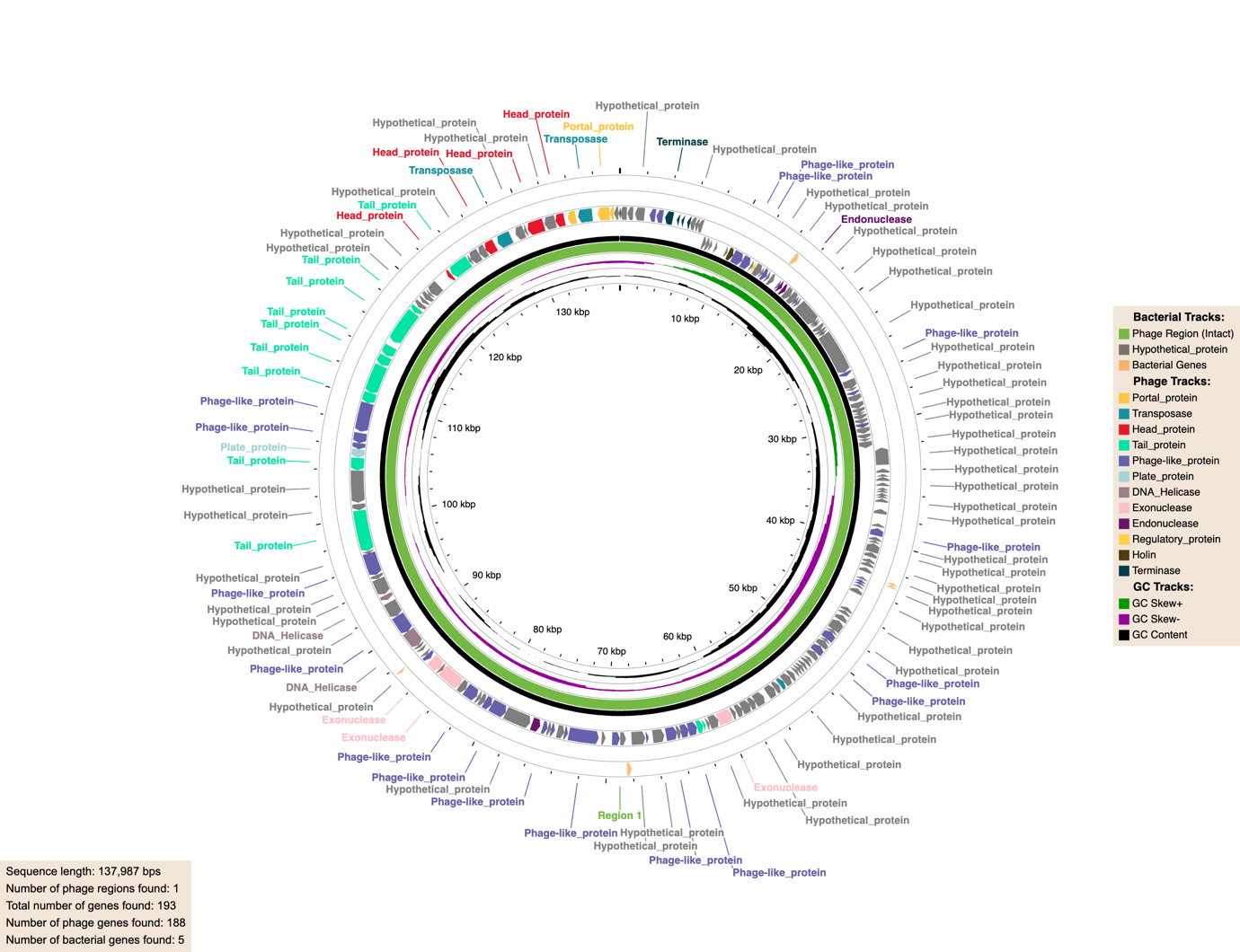

**e)**

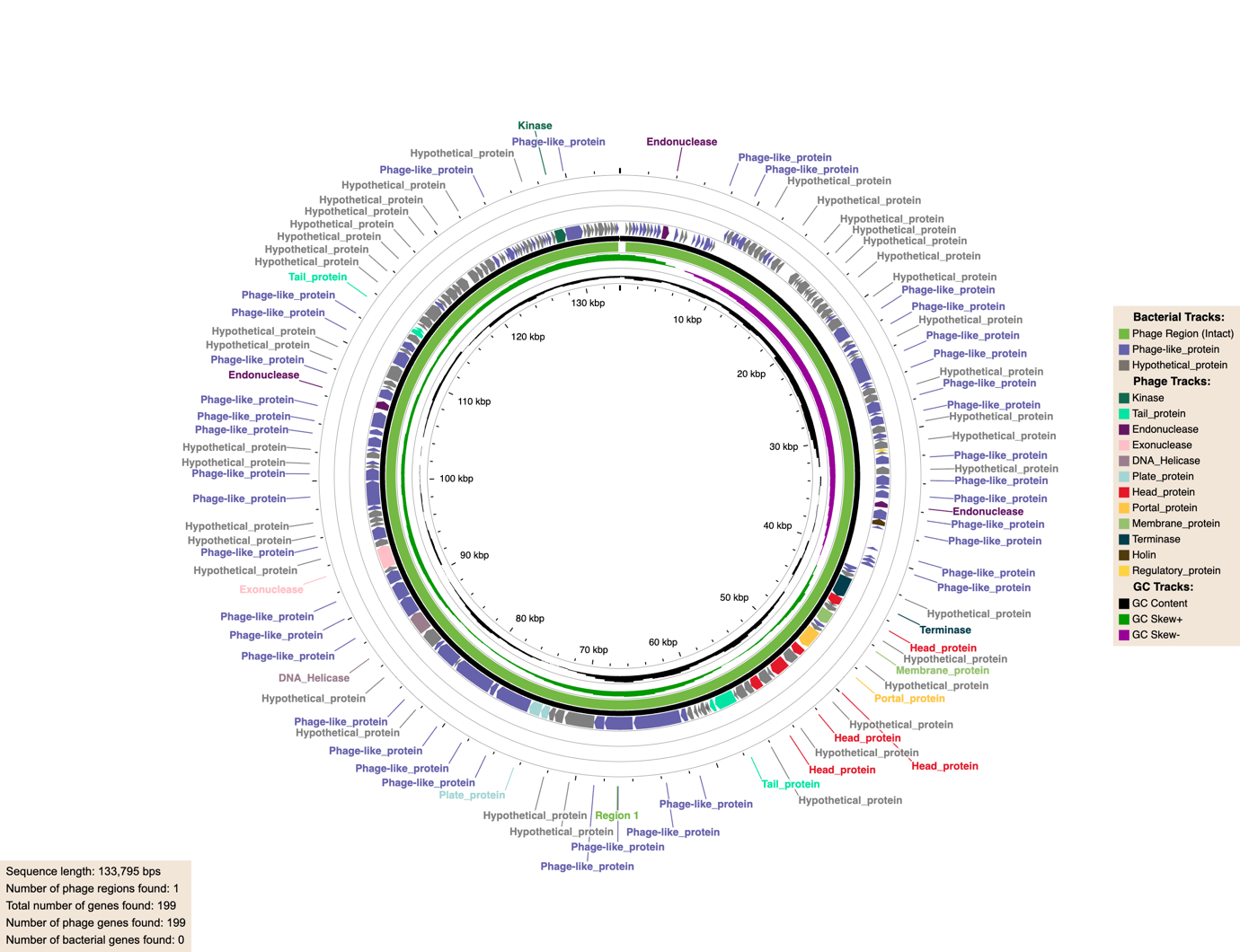

**f)**

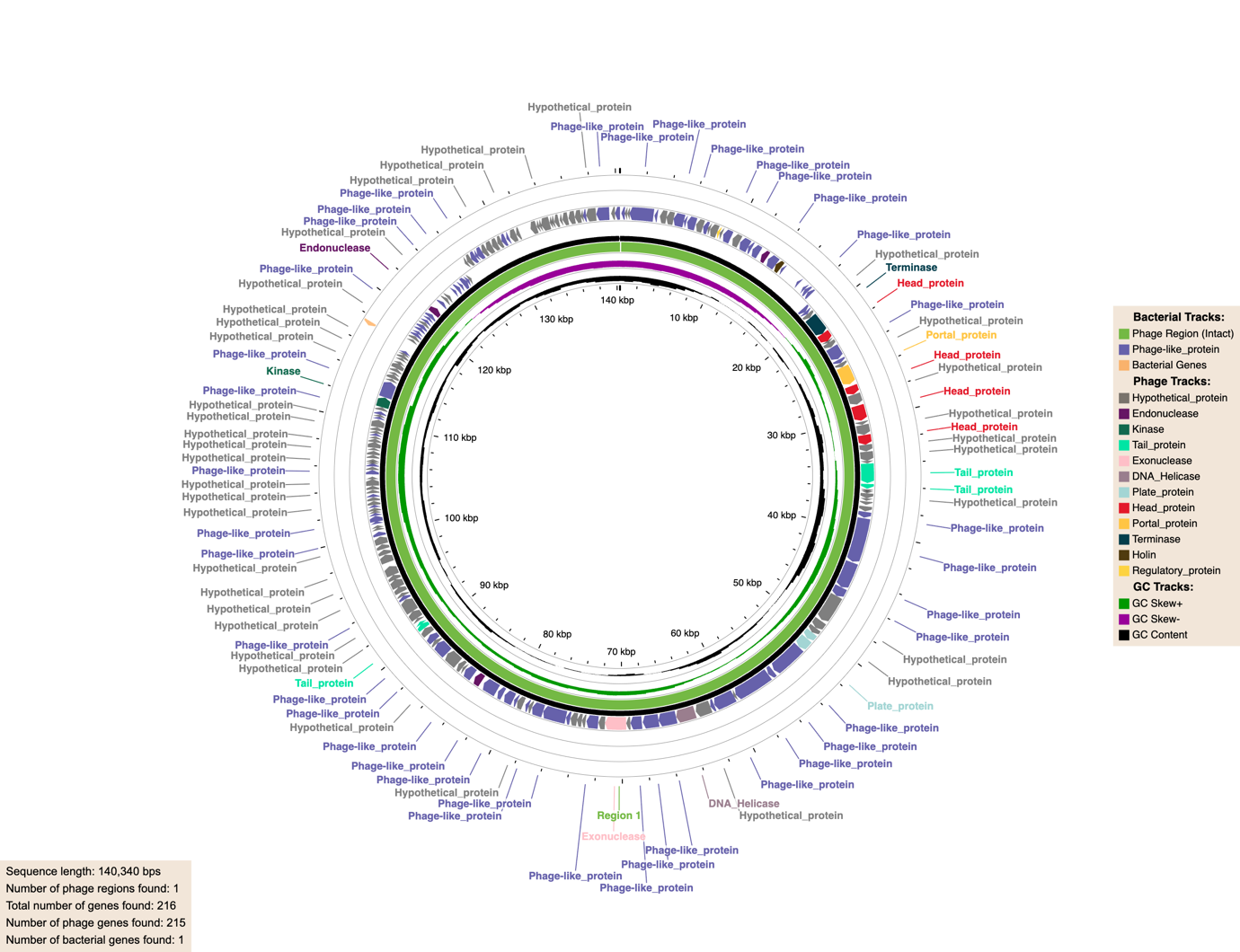

**g)**

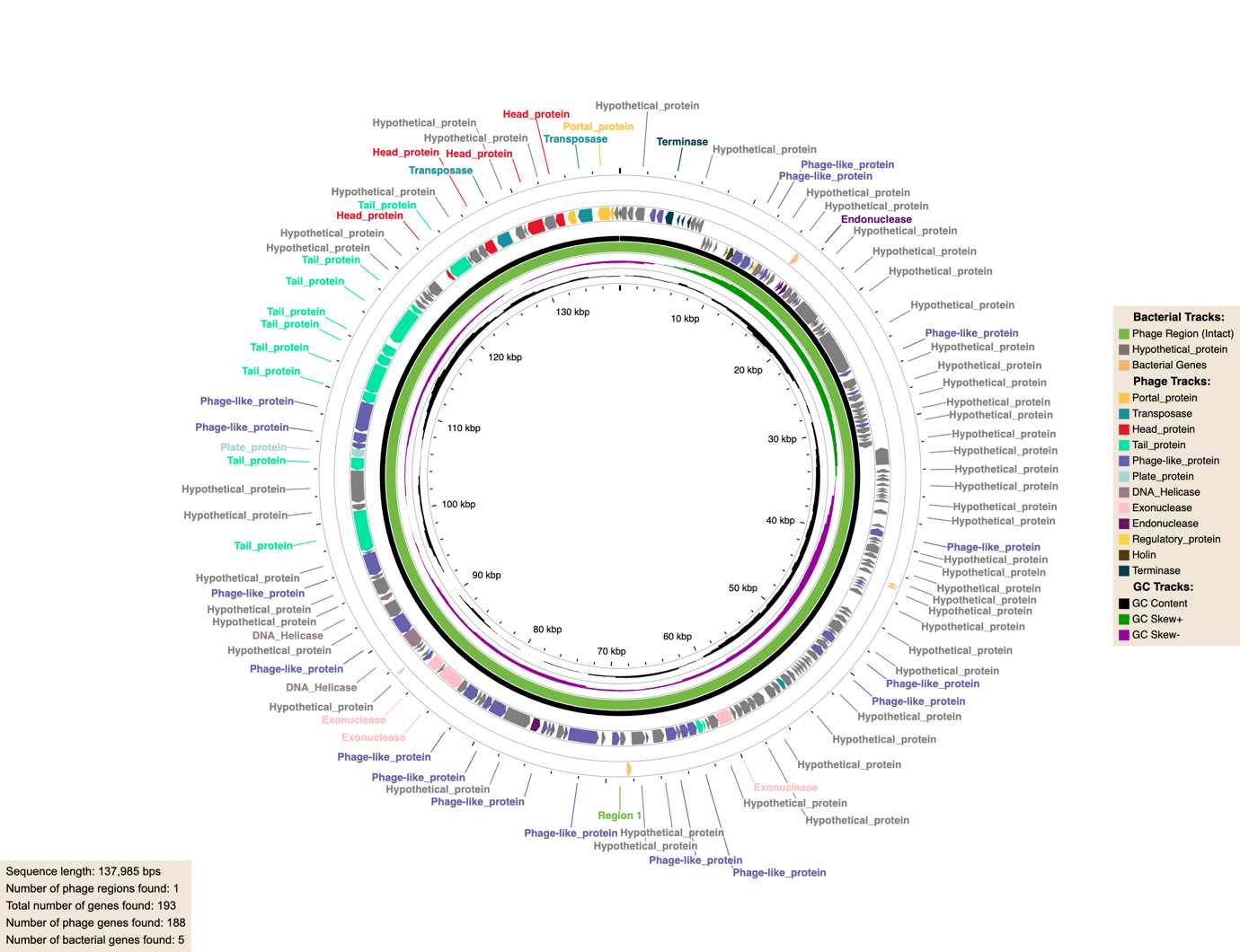

**h)**

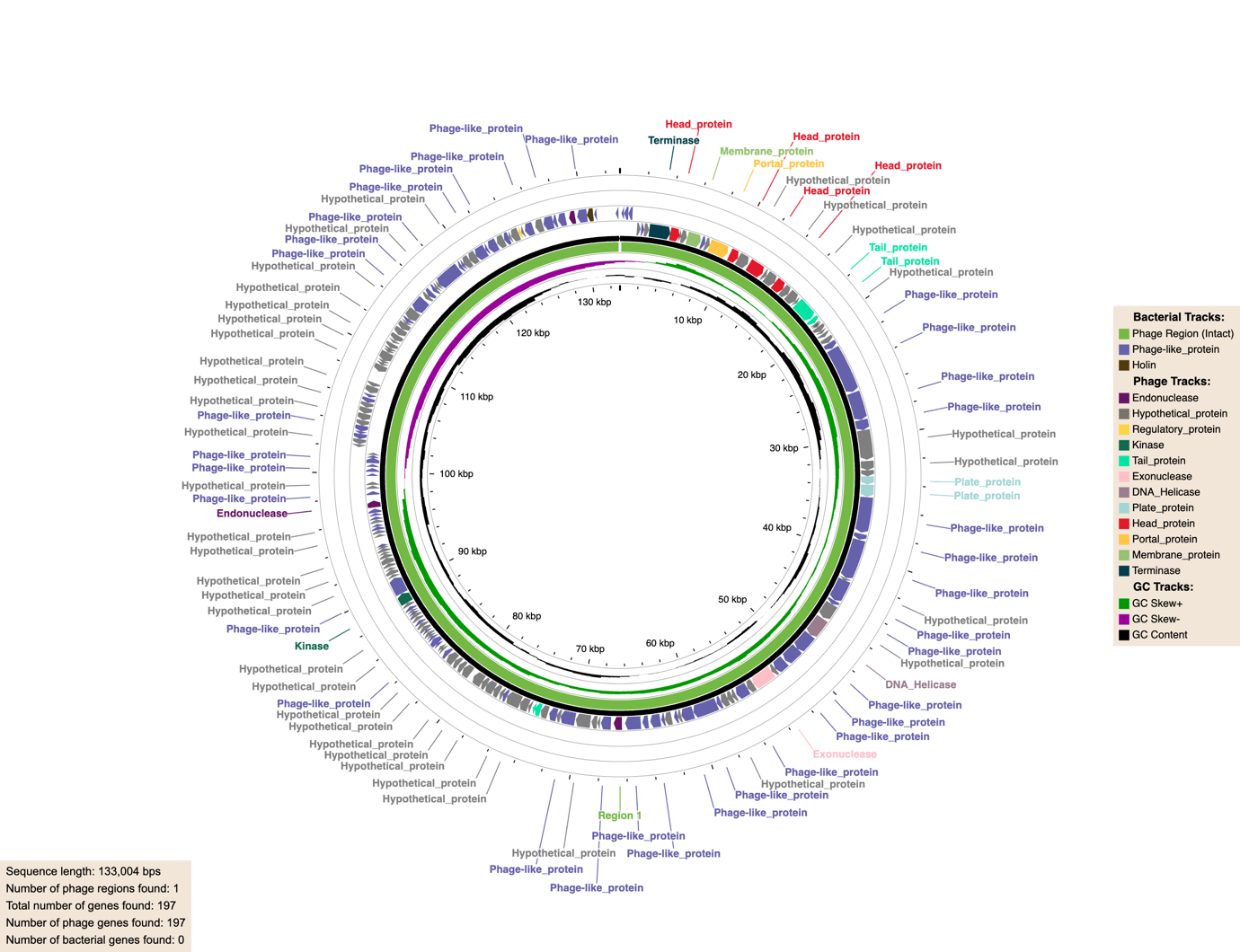

**i)**

j)
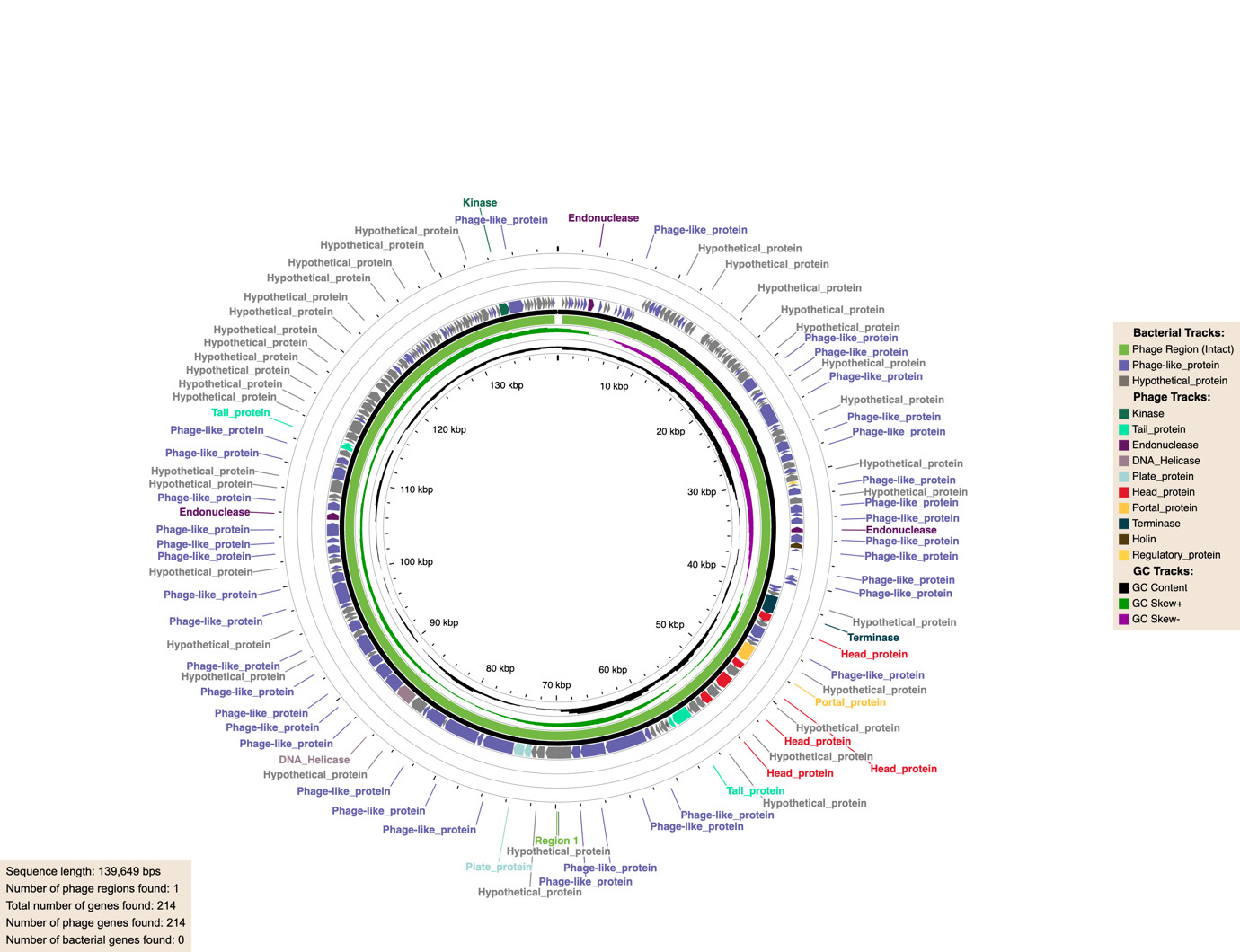

k)
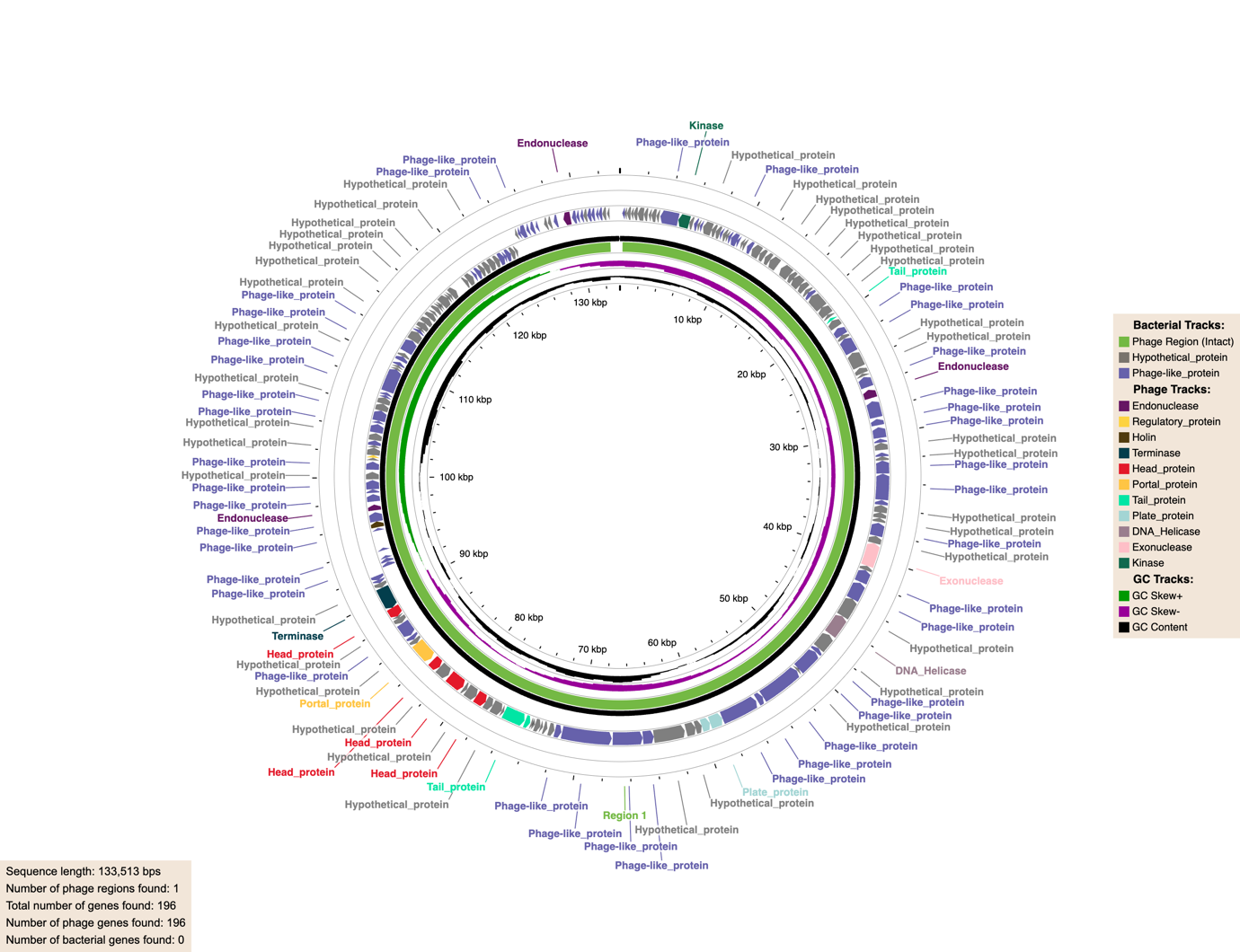

l)
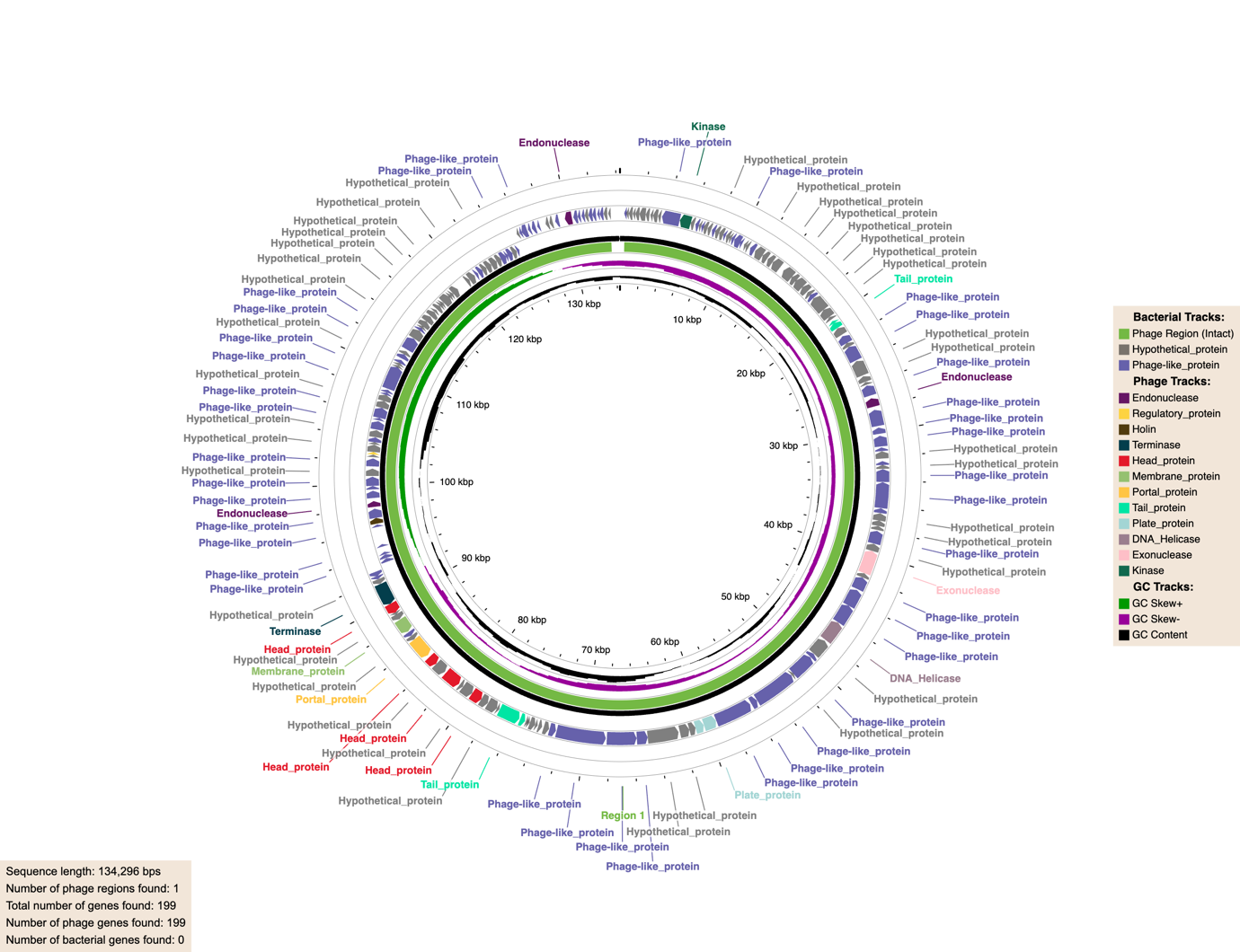

m)
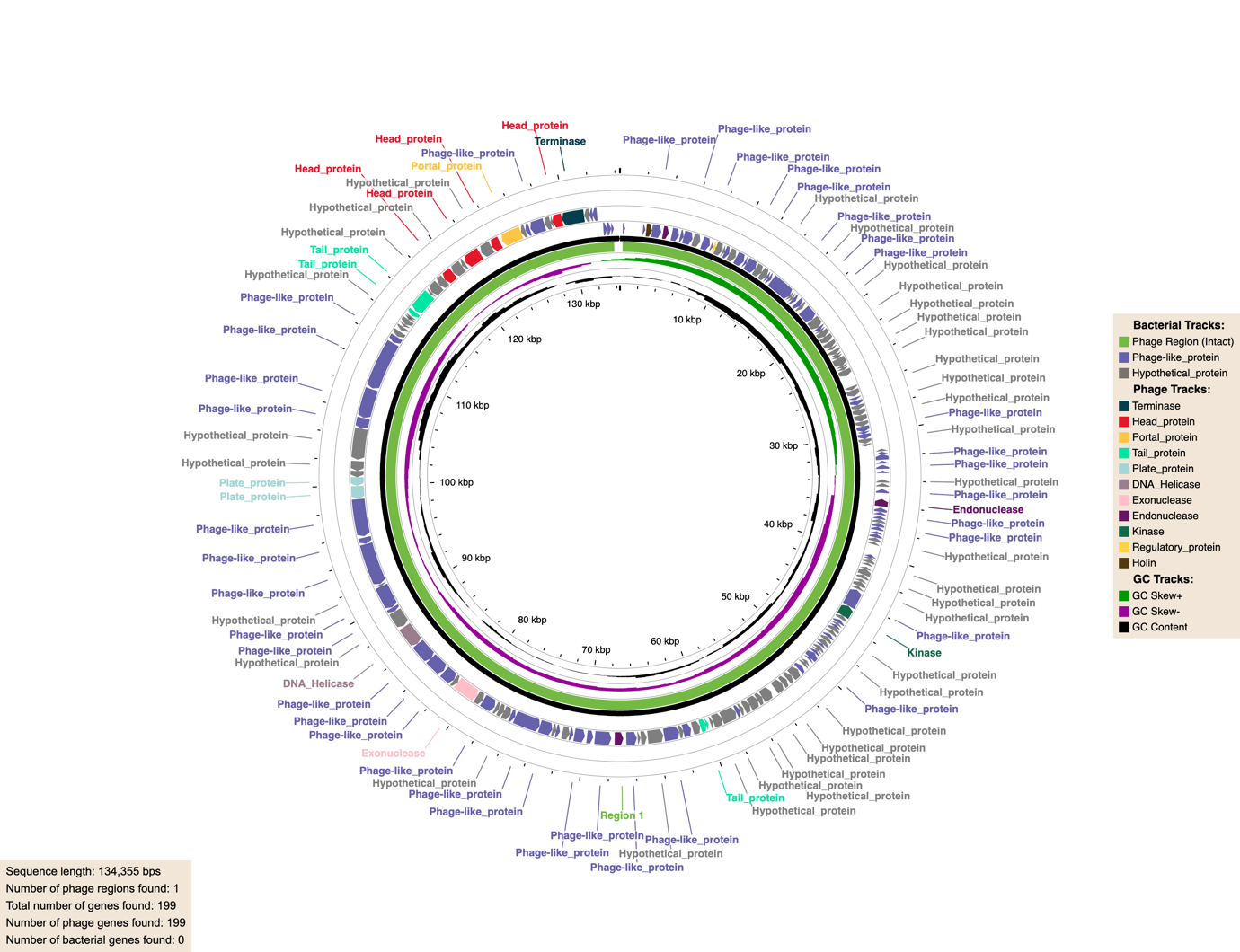

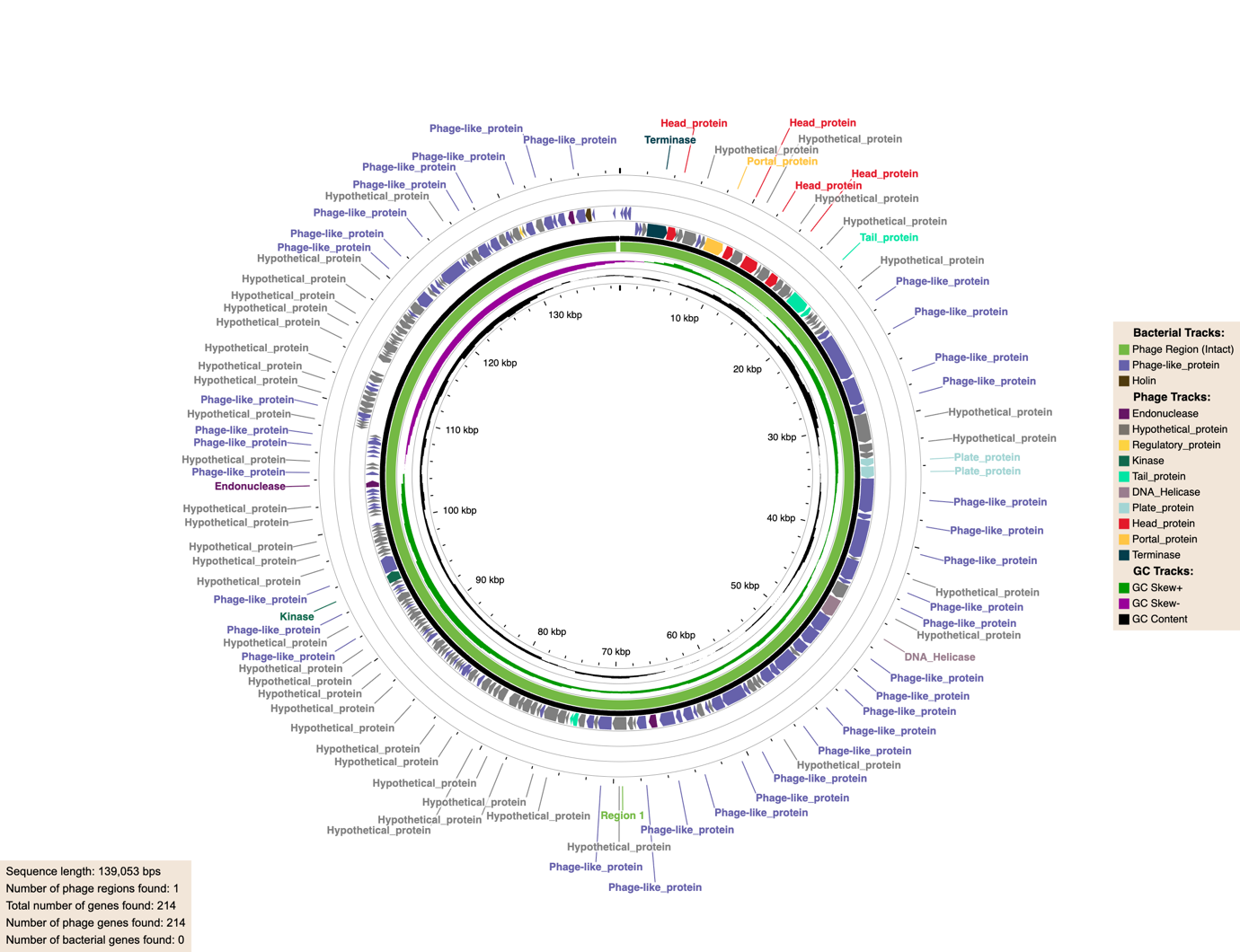

o)
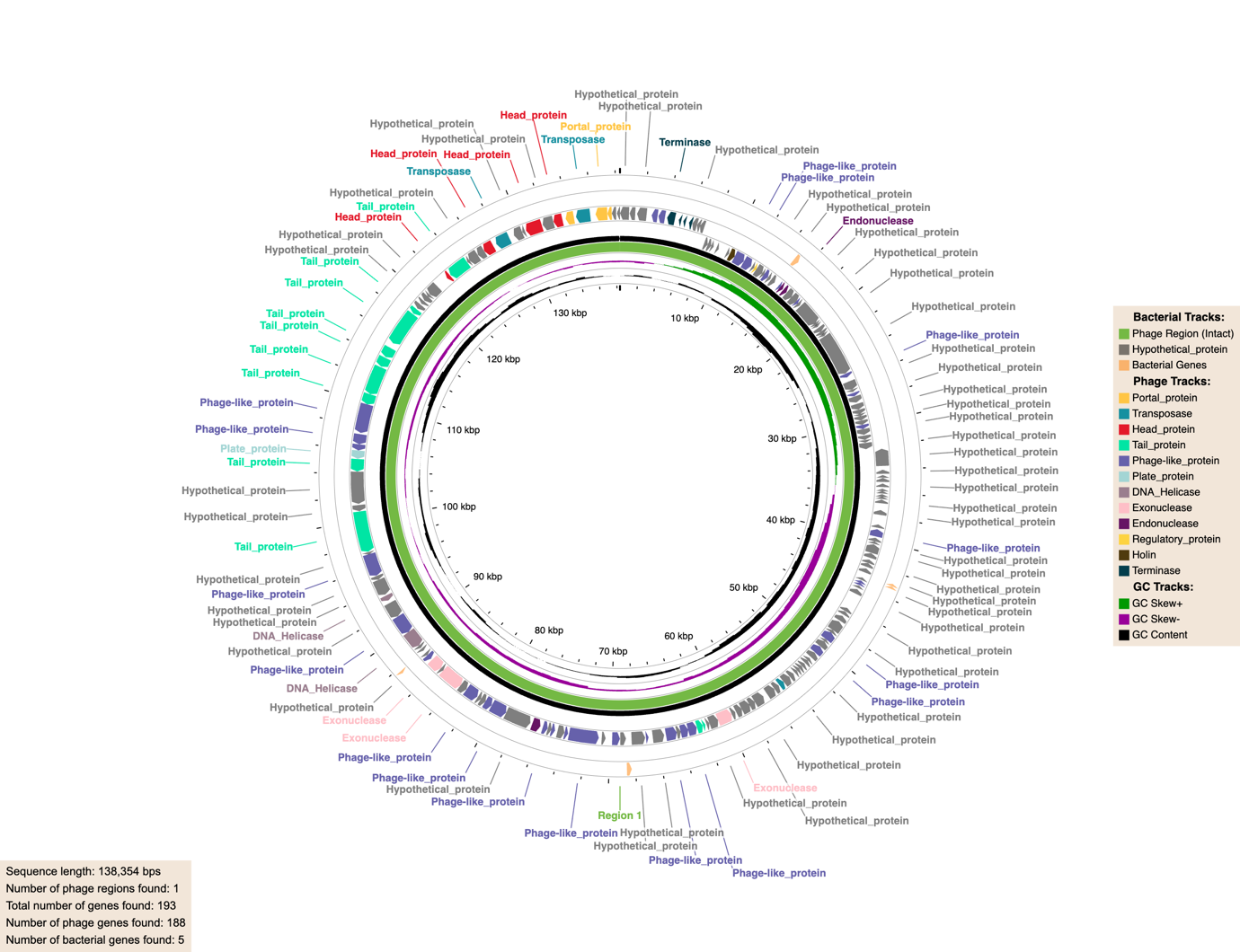
**.**

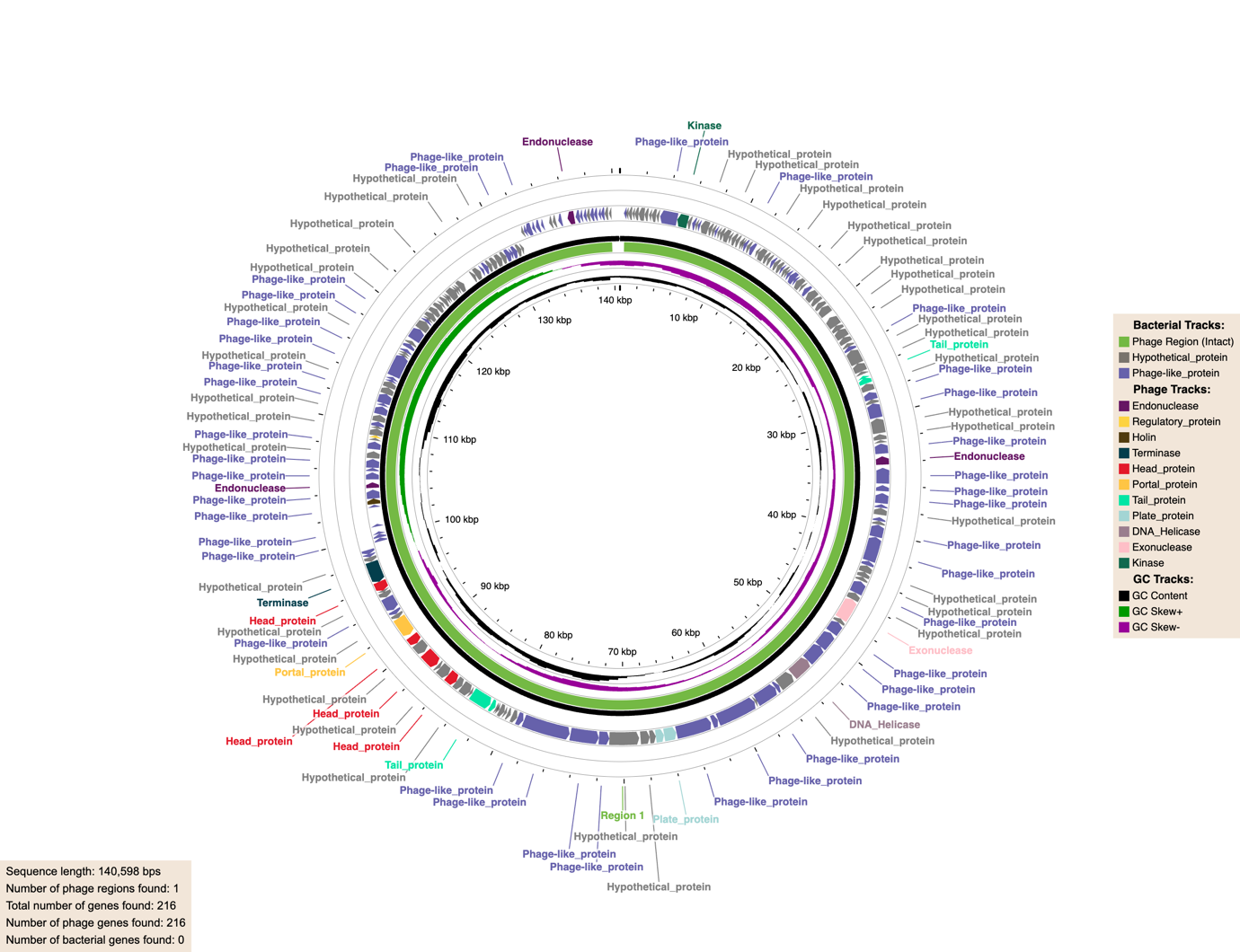

q)
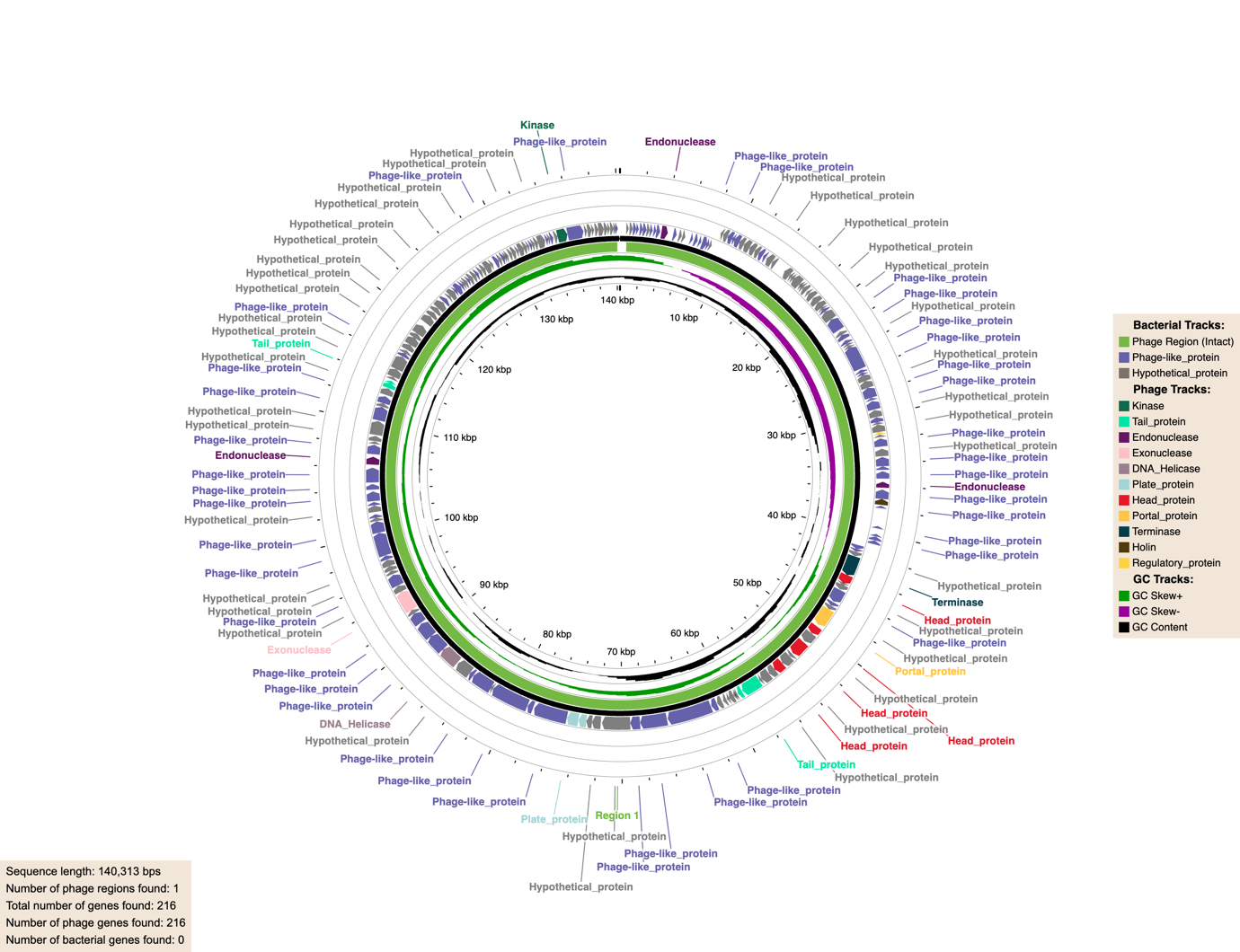

r)
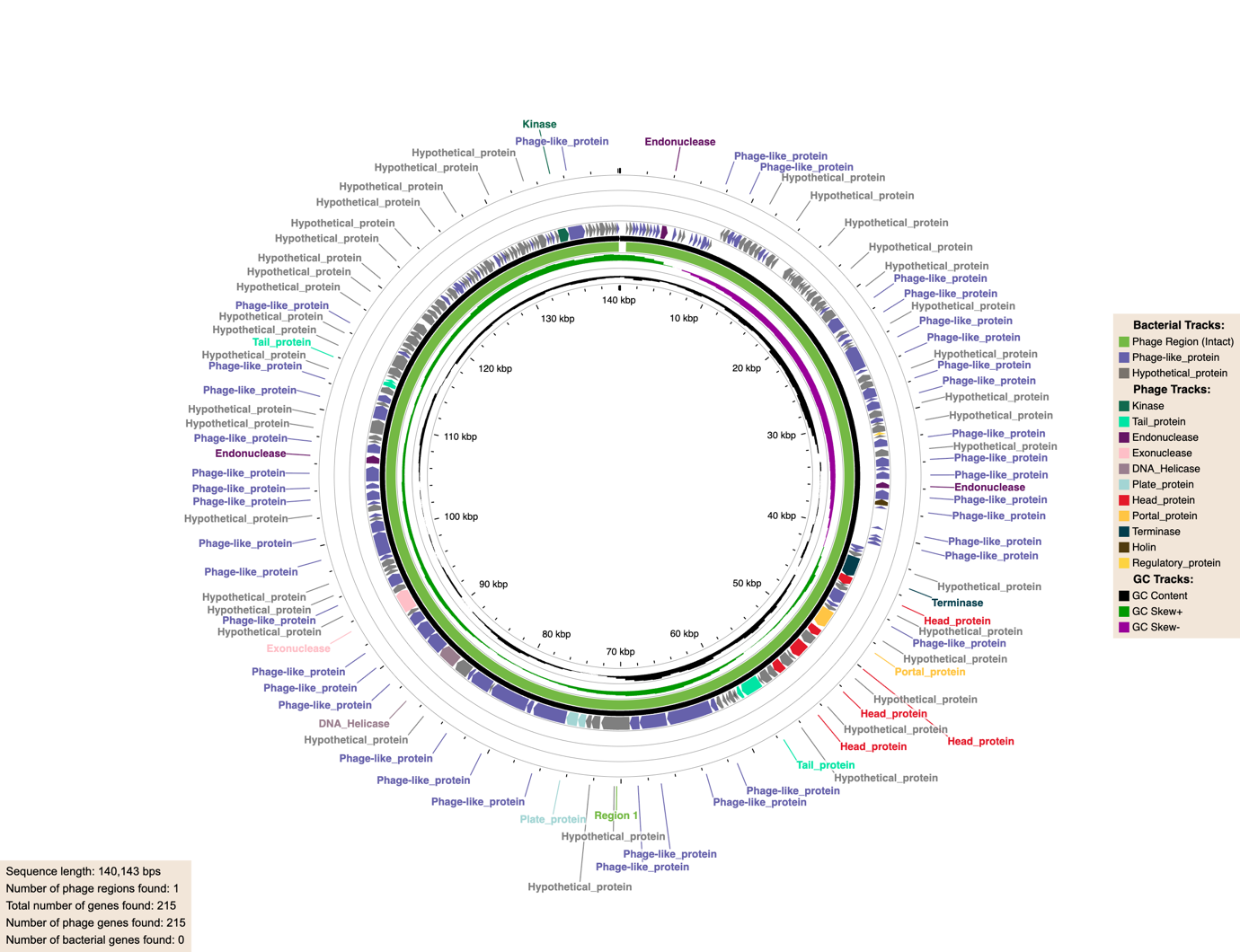

s)
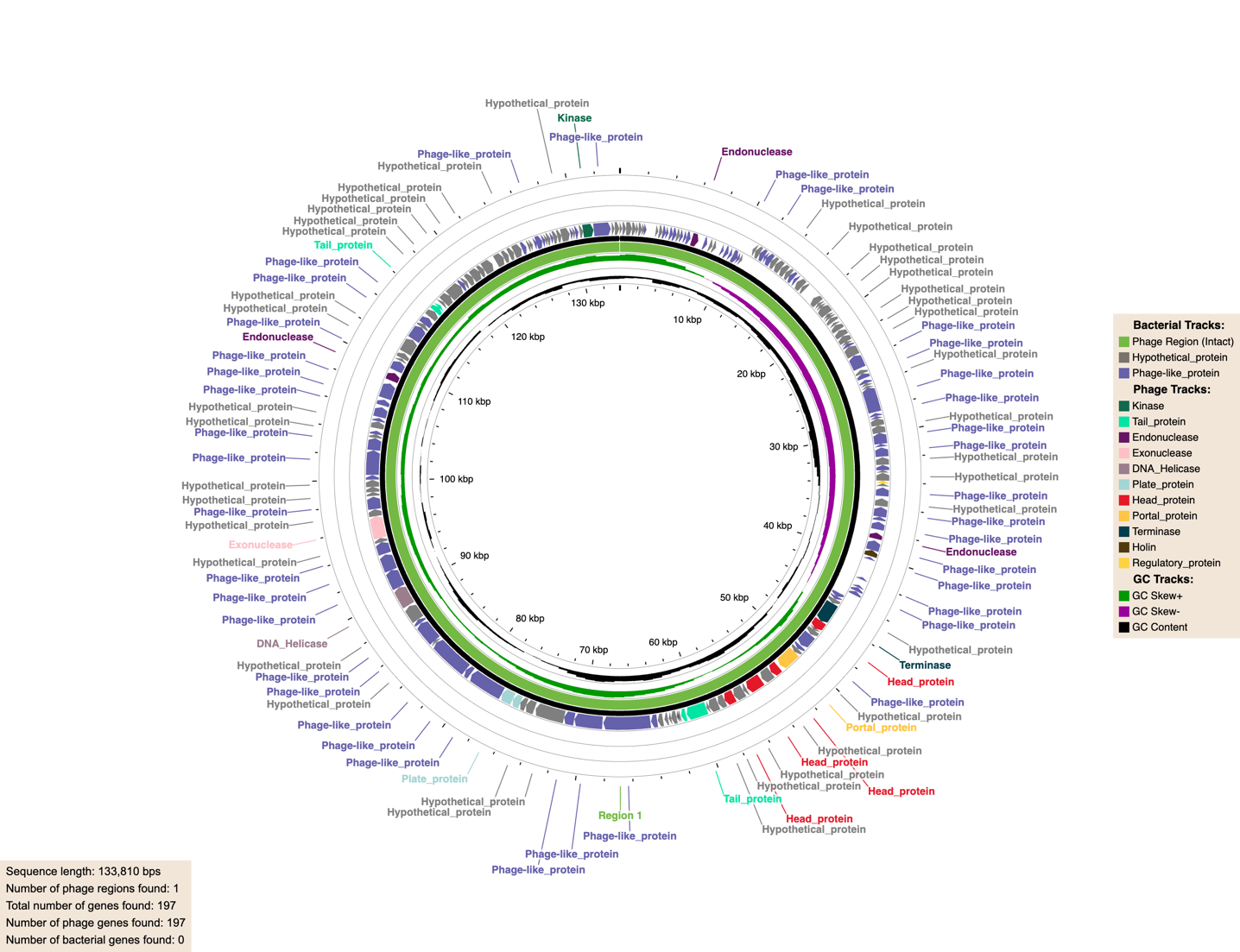

t)
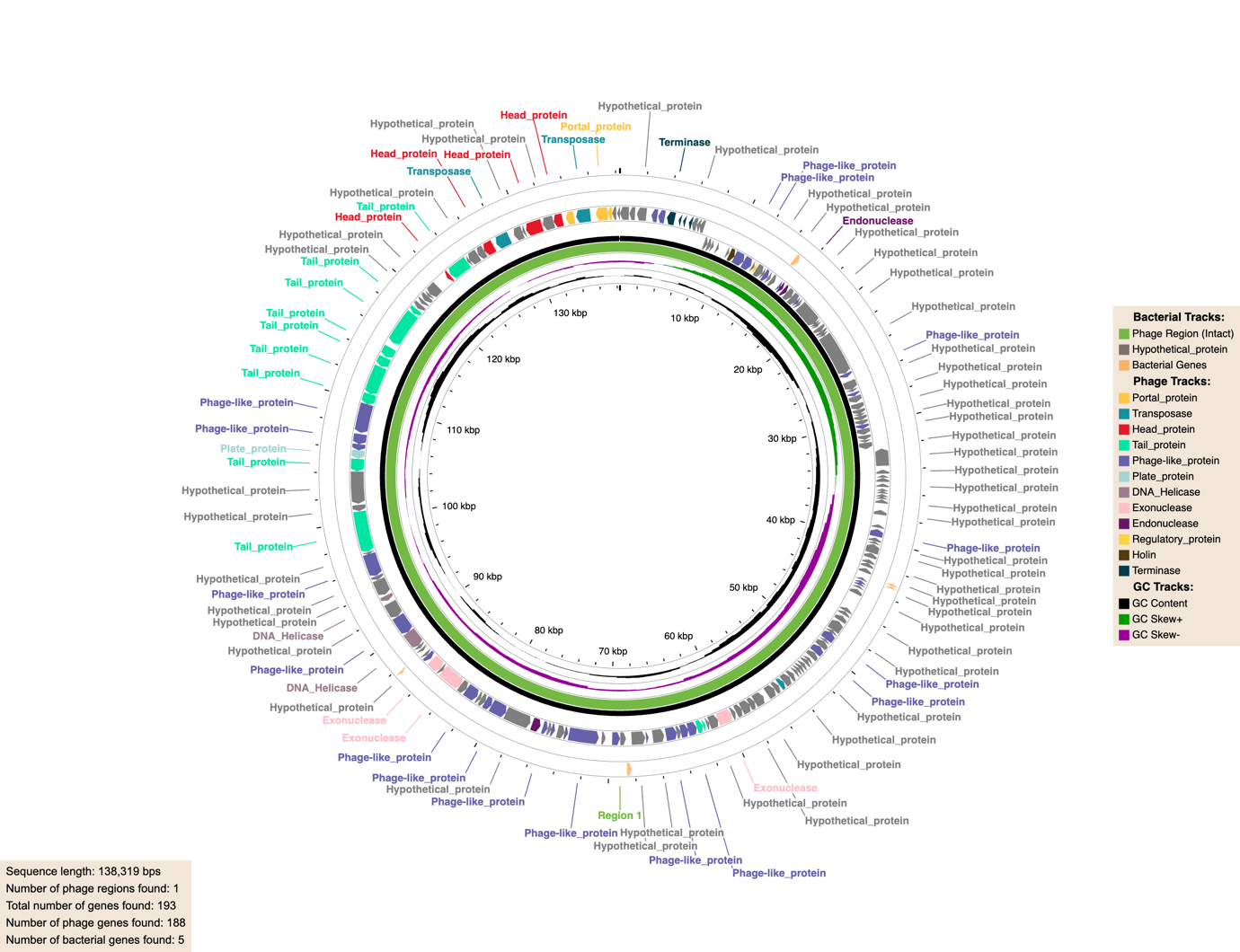

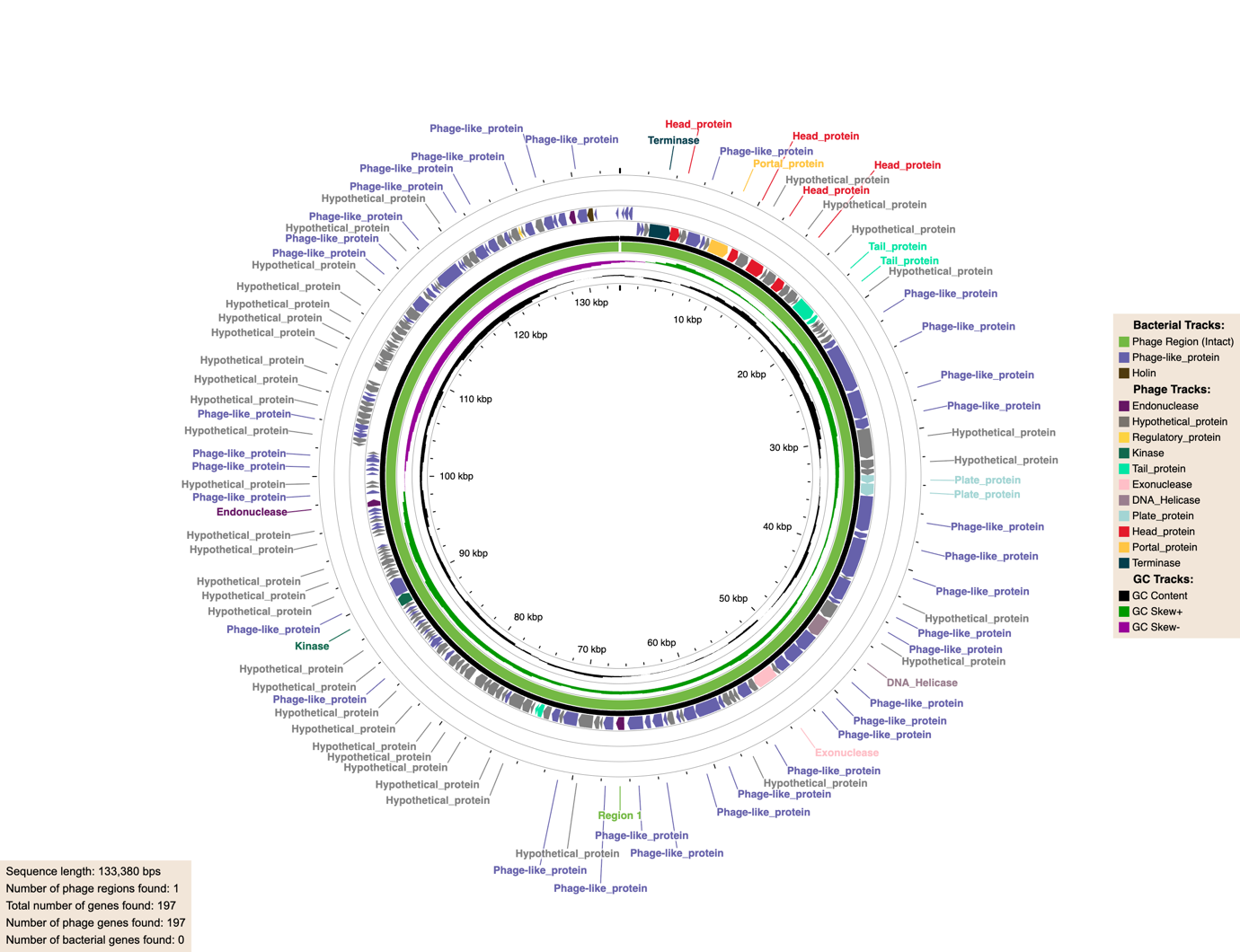

v)
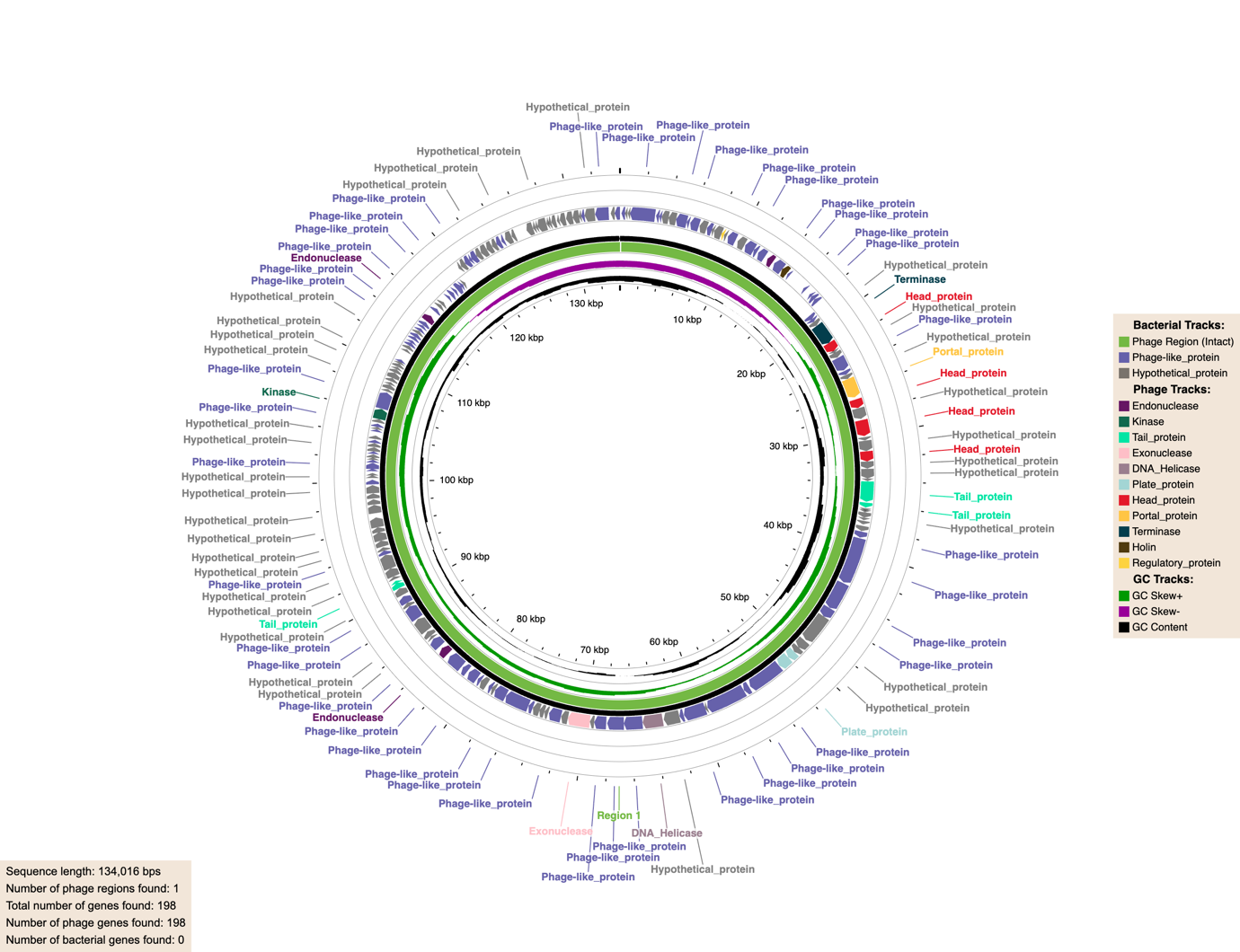

w)
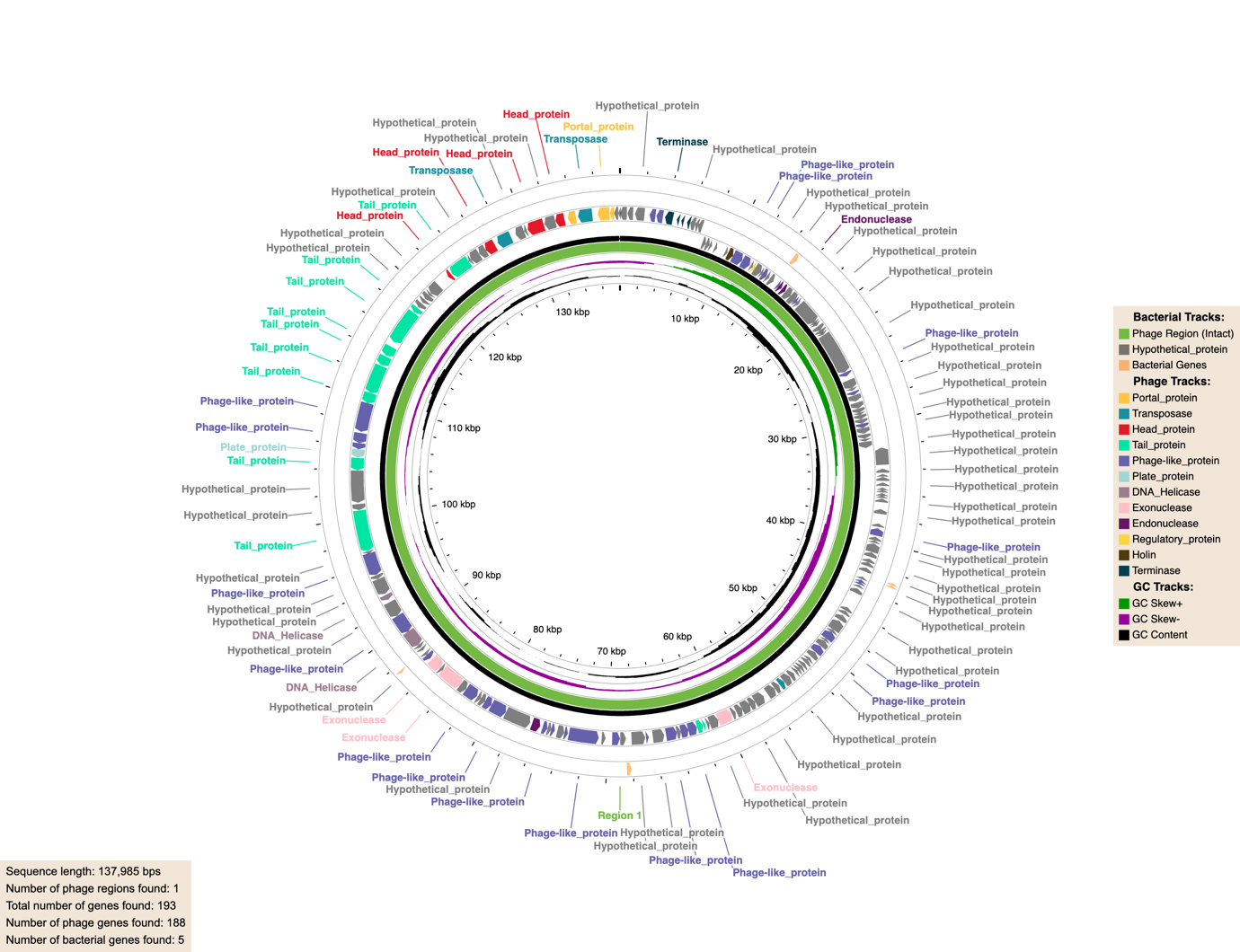

x)
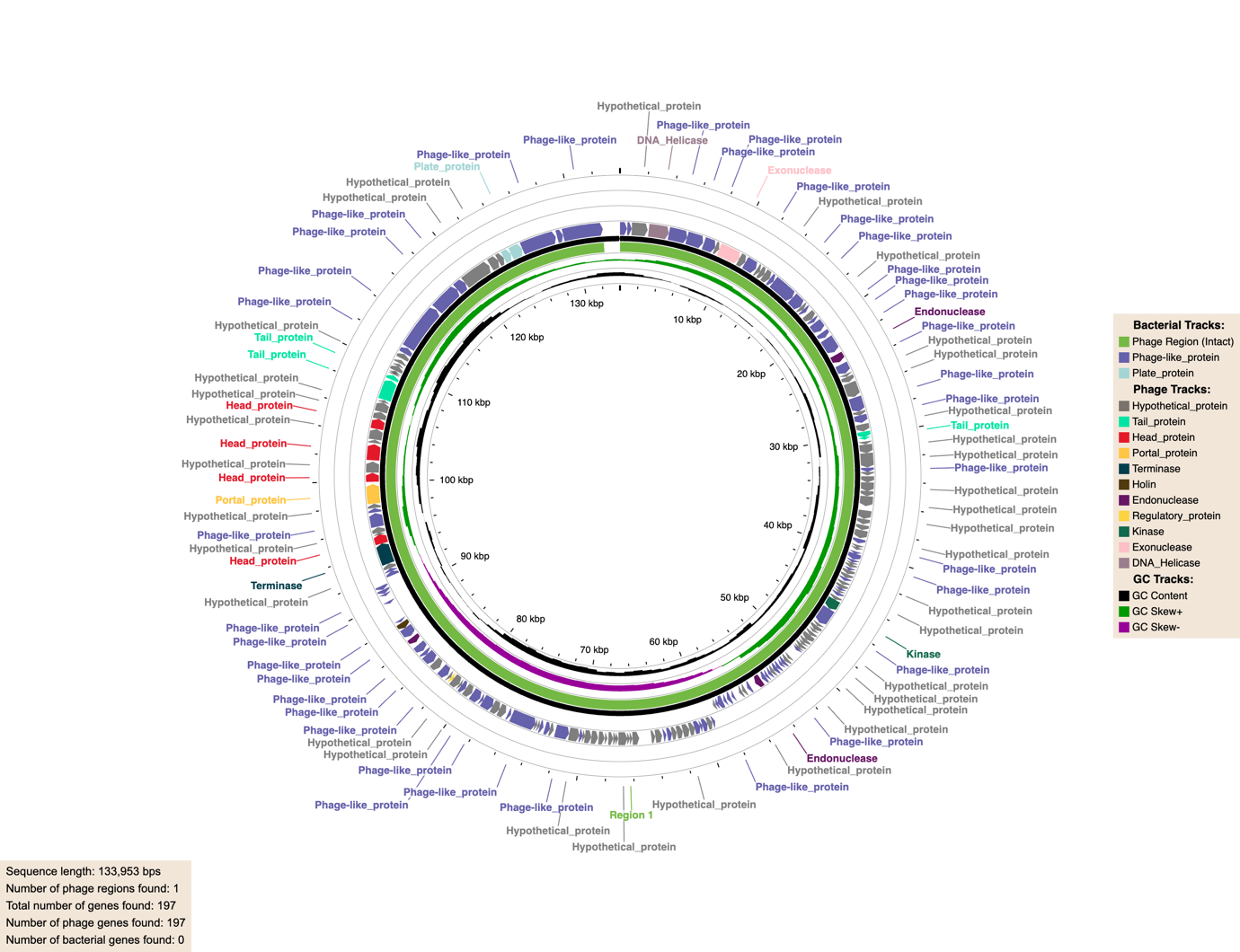

**Supplementary Figure 2: Circular genome maps of *Staphylococcus aureus* phages Φ1, Φ2, Φ6, Φ9, Φ10, Φ13, Φ17, Φ21, Φ23, Φ24, Φ26, Φ32, Φ33, Φ35, Φ36, Φ42, Φ48, Φ50, Φ53, Φ54, Φ55, Φ66, Φ68, Φ69, and *Staphylococcus epidermidis* phage Φ72.**

Circular genome representations depict the complete annotated genomes of 24 Staphylococcus aureus phages (Φ1–Φ69, selected isolates as listed) and one Staphylococcus epidermidis phage (Φ72). Genome sizes range approximately within the typical Kayvirus spectrum (~130–160 kb), with GC content conserved around ~30%, consistent with members of the family Herelleviridae. Open reading frames (ORFs) are displayed as directional arrows indicating transcriptional orientation, and are color-coded according to predicted functional categories, including structural and morphogenesis modules (capsid, portal, tail sheath, baseplate, tail fibres), DNA replication and metabolism proteins (DNA polymerase, helicase, primase, ligase), nucleotide modification and repair enzymes, host takeover factors, lysis cassette components (endolysin, holin, spanins where present), and hypothetical proteins. The modular genome architecture characteristic of lytic Kayviruses is preserved across all isolates, with conserved synteny observed in structural and replication blocks, while localised variability is evident within host-recognition and accessory gene regions, particularly in tail fibre and baseplate-associated loci that likely contribute to host-range heterogeneity. No integrase, repressor, or lysogeny-associated modules were identified, supporting their obligately lytic lifestyle. Similarly, no known AMR genes or classical virulence determinants were detected based on genome annotation and database screening. Comparative inspection of the circular maps highlights a highly conserved genomic backbone with discrete hypervariable islands potentially underpinning phenotypic differences in adsorption efficiency, burst size, and lytic activity across diverse Staphylococcus sequence types. The genome of Φ72 (*S. epidermidis* phage) exhibits overall structural conservation with the S. aureus phages while maintaining distinct variations within predicted receptor-binding modules, consistent with species-specific host adaptation. These circular representations collectively demonstrate conserved macro-architecture alongside fine-scale genetic diversity, supporting the translational potential of this phage collection for therapeutic and bioengineering applications.
